# β-lactam Antibiotics Stimulate the Pathogenicity of Methicillin-resistant *Staphylococcus aureus* Via SarA-controlled Tandem Lipoprotein Expression

**DOI:** 10.1101/353227

**Authors:** Weilong Shang, Yifan Rao, Ying Zheng, Yi Yang, Qiwen Hu, Zhen Hu, Jizhen Yuan, Huagang Peng, Kun Xiong, Li Tan, Shu Li, Junmin Zhu, Ming Li, Xiaomei Hu, Xuhu Mao, Xiancai Rao

## Abstract

Methicillin-resistant *Staphylococcus aureus* (MRSA) is a leading cause of nosocomial infections worldwide. MRSA resists nearly all β-lactam antibiotics that have a bactericidal activity and a signal inducer effect. However, studies have yet to clarify whether the inducer effect of empirically used β-lactams stimulates MRSA pathogenicity in vivo. Here, we showed that a new cluster of tandem lipoprotein genes (*tlpps*) was upregulated in MRSA in response to the subinhibitory concentrations of β-lactam induction. The increased Tlpps significantly altered immune responses by macrophages with high IL-6 and TNFα levels. The deletion of the *tlpps* mutant (N315Δ*tlpps*) significantly decreased the proinflammatory cytokine levels in vitro and in vivo. The bacterial loads of N315Δ*tlpps* in the mouse kidney were also reduced compared with those of the wild type N315. The β-lactam-treated MRSA exacerbated cutaneous infections with increased lesion size, extended illness, and flake-like abscess-formation compared with those of the nontreatment. The β-lactam antibiotics that promoted the MRSA pathogenicity were SarA dependent, and the increasing expression of *tlpps* after β-lactam treatment was directly controlled by the global regulator SarA. Overall, our findings suggested that β-lactams should be used carefully because it might lead to a worse outcome of MRSA infection than inaction in the treatment.

**Author summary:** β-lactams are widely used in practice to treat infectious diseases, however, β-lactams worsening the outcome of a certain disease is poorly understood. In this study, we have identified a new cluster of tandem lipoprotein genes (*tlpps*) that is upregulated in the major clinically prevalent MRSA clones in response to the subinhibitory concentrations of β-lactams induction. The major highlight in this work is that β-lactams induce SarA expression, and then SarA directly binds to the *tlpp* cluster promoter region and upregulates the *tlpp* expression in MRSA. Moreover, the β-lactam stimulated Tlpps are important virulence factors that enhance MRSA pathogenicity. The deletion of the *tlpps* mutant significantly decreases the proinflammatory cytokine levels in vitro and in vivo. The β-lactam induced Tlpps enhance the host inflammatory responses by triggering the expression of IL-6 and TNFα, thereby promoting bacterial colonization and abscess formation. These data elucidate that β-lactams can worsen the outcome of MRSA infection through the induction of *tlpps* that are controlled by the global regulator SarA.

## Introduction

Infectious diseases caused by bacteria are common and adversely affect human health worldwide. The discovery of antibiotics for antibacterial application was a remarkable achievement in the 20th century. In the therapeutic use of antibiotics in humans and animals, bacteria encounter wide gradients of antibiotic concentrations in host bodies [1]. Antibiotics can serve as signal inducers, in addition to their clinically important antibacterial activity, and influence the physiological characteristics of bacteria and trigger various cellular responses in bacterial species. Low levels of antibiotics can induce extracellular DNA release, virulence factor production, and biofilm formation [2,3], resulting in a worse outcome of bacterial infections. Therefore, the mechanisms underlying the stimulation of pathogenicity in a bacterial population at subinhibitory antibiotic concentrations should be understood.

Methicillin-resistant *Staphylococcus aureus* (MRSA) is a leading pathogen with notable pathogenic effects. MRSA causes a wide range of diseases, including acute skin and soft tissue infections, chronic and persistent endocarditis, osteomyelitis, and pneumonia [4,5]. MRSA infections cause greater morbidity and mortality than methicillin-susceptible *S. aureus* (MSSA) infections do [6,7]. However, the underlying mechanisms of these effects remain unclear. Several studies have suggested that as-yet-unidentified virulence factors or inappropriate treatments contribute to the poor outcome of MRSA infections [6,8,9]. It has been reported that between 30% and 80% of individuals infected with MRSA are inappropriately treated often with β-lactam antibiotics because of the failure to recognize MRSA infection initially [10,11]. Accumulated data have demonstrated that the subinhibitory concentrations of β-lactam antibiotics can promote *S. aureus* pathogenicity by increasing the expression of alpha-toxin [12], Panton-Valentine leukocidin (PVL) [2], enterotoxins [13], or staphylococcal protein A (SPA) in vitro [14]. Nevertheless, the contribution of the certain altered virulence factor to the MRSA pathogenesis in vivo has yet determined, and the molecular mechanisms underlying β-lactams modulating MRSA pathogenesis remain largely unknown.

Lipoproteins (Lpps) are an abundant family of proteins anchored in the bacterial membrane, and they account for at least 2% of a bacterial proteome [15,16]. *S. aureus* encodes 55-70 putative Lpps, and approximately 50% of these Lpps are annotated as chaperones or as transporters for amino acids, peptides, iron, and zinc [17]. Many of the proposed Lpps (more than 30%) in *S. aureus* are conserved hypothetical proteins of unknown functions [16]. Most virulent MRSA strains, such as USA300, carry a conserved genomic island termed vSaα (nonphage and nonstaphylococcal cassette chromosome genomic island) that encodes numerous homologous *lpps* arranged in tandem, which is referred to as “tandem lipoproteins” (*tlpps*) or “lipoprotein-like” (*lpl*) [15,18]. This *tlpp* cluster likely represents the paralogous genes that have diverged after a duplication event in *S. aureus* [17]. MRSA USA300 belonging to the clonal complex CC8 carries 15 (22%) hypothetical Tlpps. Of these Tlpps, 9 are specific to the vSaα genomic island [15]. By comparison, N315 belonging to the clonal complex CC5 carries 12 (21%) hypothetical Tlpps. Of these Tlpps, 9 Lpl proteins are specific to the vSaα genomic island (S1 Table).

Some staphylococcal Lpps can trigger host cell invasion, increase bacterial pathogenicity, and contribute to the epidemic of CC8 and CC5 strains [19,20]. However, the exact roles of the Tlpp proteins are unclear. Whether these Tlpp proteins can be induced by the subinhibitory concentrations of antibiotics and contribute to the pathogenesis of MRSA have yet to be determined. In this study, we demonstrated that a new *tlpp* cluster of MRSA was upregulated in response to the subinhibitory concentrations of β-lactam induction. The increased Tlpps in MRSA significantly altered immune responses by increasing the IL-6 and TNFα levels of the macrophages. Null Tlpp MRSA mutant infection decreased the IL-6 and TNFα levels in serum and the bacterial burdens in kidney of a mouse model. β-lactam-treated MRSA N315 exhibited an increased pathogenicity with severe cutaneous infection and abscess formation. Moreover, the β-lactam antibiotics that promoted pathogenicity in MRSA were SarA dependent, and the increasing Tlpp expression after β-lactam treatment was directly controlled by the global regulator SarA.

## Results

### β-lactam antibiotics stimulated tandem lipoprotein expression in MRSA

It is reported that the subinhibitory concentrations of β-lactam antibiotics can induce the production of some *S. aureus* toxins, such as alpha-toxin [12], PVL [2], and enterotoxins [13], or immune evasion molecules, such as SPA [14]. Here, *S. aureus* N315, which is a globally prevalent sequence type 5 (ST5) MRSA strain [21], was tested for its antibiotic response to identify new factors contributing to MRSA pathogenesis in antibiotic induction. The minimal inhibitory concentrations (MICs) of β-lactam antibiotics, including oxacillin (OXA), methicillin (MET), cefoxitin (FOX), imipenem (IMI), meropenem (MER), chloramphenicol (CHL), vancomycin (VAN), kanamycin (KAN), and erythromycin (ERY) against N315 were determined (S2 Table). The sodium dodecyl sulfate-polyacrylamide gel electrophoresis (SDS-PAGE) revealed that a protein band (approximately 30 kDa) was upregulated in the subinhibitory concentrations of β-lactam-treated bacteria compared with the nontreatment control (Fig 1A). By contrast, CHL, VAN, KAN, and ERY did not have induction effects on this protein band. Further observations indicated that the subinhibitory concentrations of OXA exerted a broad-spectrum induction effect on other major clinically prevalent MRSA clones, including ST88, ST239, ST59, ST1, and ST398, which displayed the same upregulated protein band as that in the OXA-treated MRSA compared with the nontreatment one (Fig 1B).

**Fig 1.**
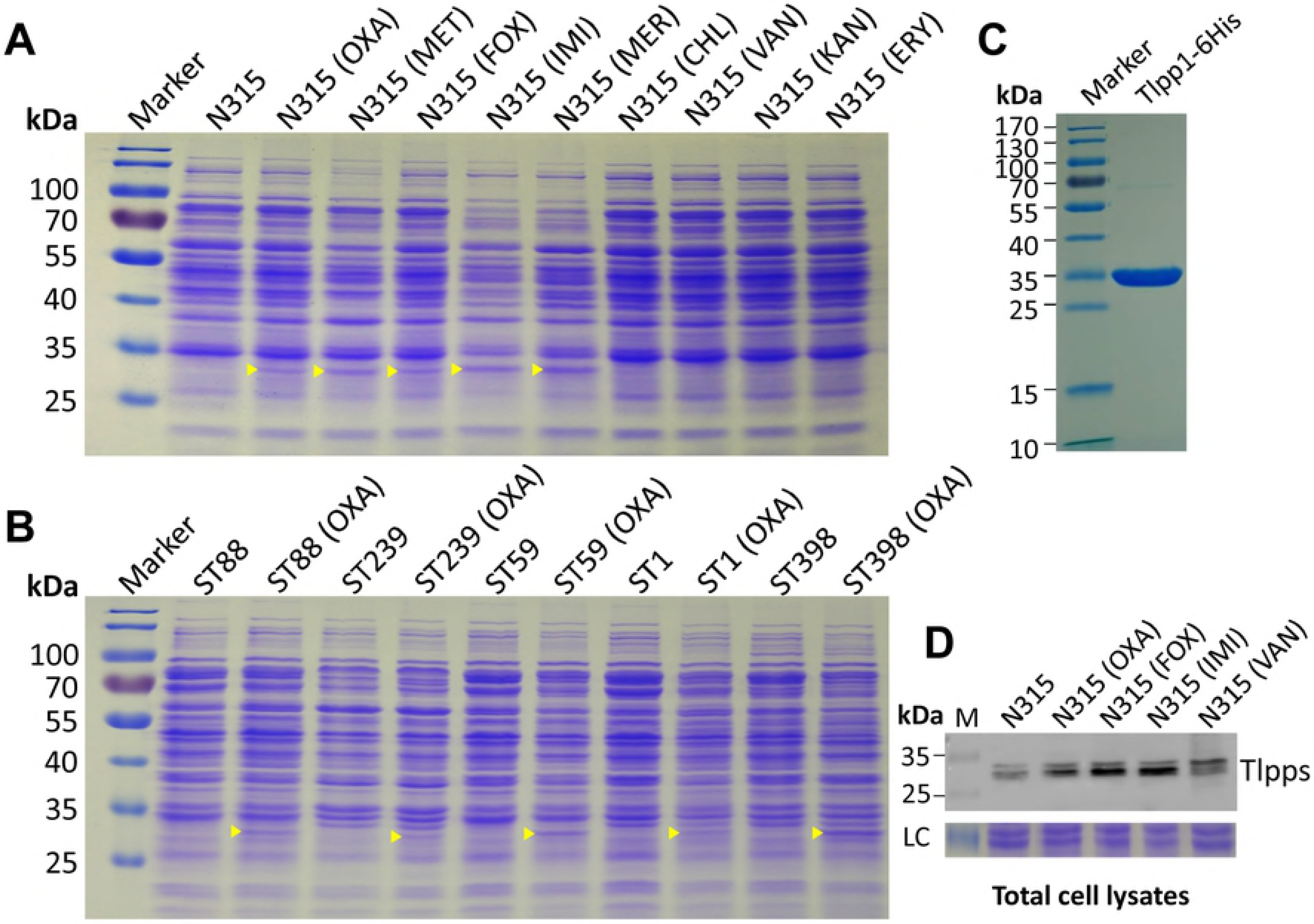
Tlpps upregulated in MRSA post-treated with the subinhibitory concentrations of antibiotics. (A) The proteins of different antibiotic-induced MRSA N315 were separated through SDS-PAGE and stained with Coomassie brilliant blue. An untreated N315 served as the negative control, and the molecular weights of the protein marker were indicated on the left. The upregulated protein bands upon β-lactam antibiotic treatment were denoted by yellow triangles. (B) Other major clinically prevalent MRSA strains represented by sequence type (ST) were cultured in the absence or presence of OXA (2 μg/ml, unless specifically stated, S2 Table and S9 Fig). SDS-PAGE was then performed. The yellow triangles indicated that certain upregulated protein bands were similar to those of β-lactam-induced MRSA N315 (ST5). (C) SDS-PAGE analysis of the recombinant Tlpp1 proteins. (D) Western blot analysis of β-lactam-induced proteins in N315. The full-length blot was presented in S10 Fig.

The protein band was excised from the SDS-PAGE gel and analyzed through liquid chromatography tandem mass spectrometry (LC-MS/MS) to characterize the β-lactam-induced proteins in the MRSA strains. The detected peptides matched with 68 proteins in the N315 proteome (S3 Table). Most known metabolic enzymes were excluded, and three putative Tlpps, namely, Tlpp3 (SA2273), Tlpp2 (SA2274), and Tlpp1(SA2275), encoded by a consecutive gene cluster were selected on the basis of theoretical molecular weights for the analysis (Fig 2A and S1 Table). SA2273, SA2274, and SA2275 were annotated as hypothetical proteins in the N315 genome (GenBank accession no. BA000018.3). A typical Lpp precursor contained a signal peptide at the N-terminal and a characteristic conserved three-amino acid lipobox in front of the invariable cysteine [(LVI) (ASTG) (GA)↓**C**] [16,17]. Both Tlpp1 and Tlpp3 possessed signal peptides and “lipobox” sequences, whereas Tlpp2 comprised a transmembrane helix domain at the N-terminal (S1A Fig). These proteins were annotated as Tlpps belonging to a domain of unknown function (DUF) 576 protein family on the Pfam database [22]. Considering that Tlpps account for more than 62.6% of amino acid identity (S1B Fig), we prepared recombinant Tlpp1 proteins (Fig 1C) and the corresponding antibodies, and verified the authority of the β-lactam-stimulated proteins through Western blot analysis (Fig 1D). Although several MRSA clones, such as ST8 and ST239, contained only two of the three *tlpps* in their genomes, the *tlpp* cluster was widely distributed among the major prevalent MRSA clones (S4 Table). This observation was consistent with the finding that several major MRSA clonal strains upregulated Tlpps in response to the subinhibitory concentrations of β-lactam treatment (Fig 1B).

**Fig 2.**
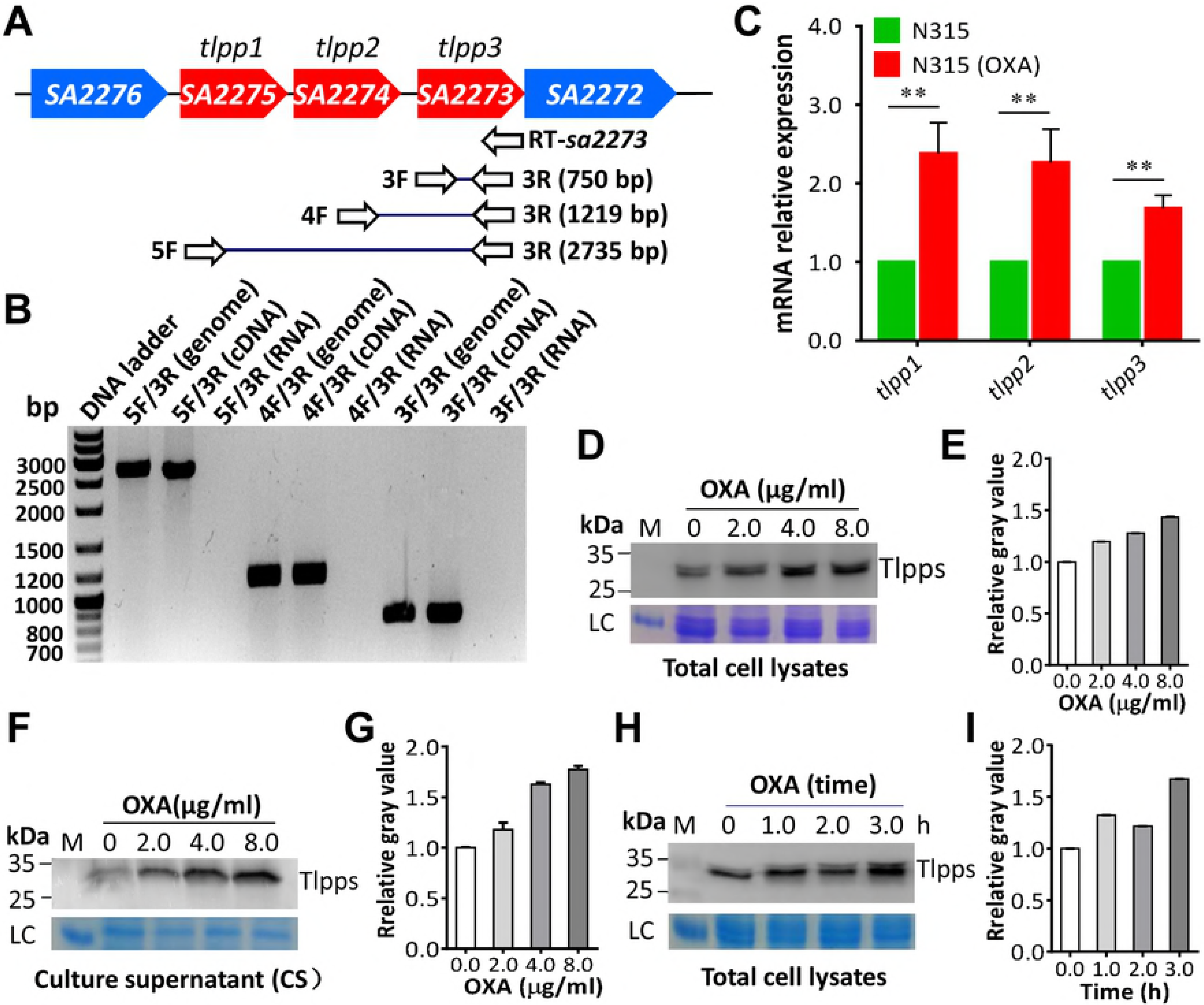
Tlpp genes were co-transcribed and upregulated by OXA in a dose- and time-dependent manner. (A) Genetic organization of the *tlpp* cluster in MRSA N315 genome and the location of primers designed for RT-PCR. (B) *tlpp1, tlpp2*, and *tlpp3* were co-transcribed tested by RT-PCR. Agarose gel electrophoresis analysis of PCR products amplified with the genomic DNA, reverse-transcribed cDNA, and RNA as templates. (C) RT-qPCR detection of the expression level of *tlpp1, tlpp2,* and *tlpp3* in N315 with or without OXA treatment was administered. ** *P* < 0.01. (D) Western blot analysis of the Tlpp levels in N315 total cell lysates after different concentrations of OXA treatment was administered. The molecular weights of the protein marker (M) were indicated on the left. LC, loading control. (E) Evaluation of gray value of the N315 Tlpps in each lane using ImageJ software. The relative value of Tlpps/loading control (LC) in the first lane (0 μg/ml OXA) was adjusted to 1.0, and the relative gray values in other lanes were calculated and indicated. (F) Western blot analysis of the Tlpp levels in N315 culture supernatant (CS) after different concentrations of OXA treatment was administered. (G) Evaluation of gray value of Tlpps in N315 culture supernatant. (H) Western blot analysis of the Tlpp levels in N315 after OXA treatment was administered with time indicated. (I) Evaluation of gray value of Tlpps in N315 after OXA treatment was given with time indicated.

### *tlppl, tlpp2,* and *tlpp3* were co-transcribed, and their expression was induced by β-lactam antibiotics in a dose- and time-dependent manner

Reverse-transcription polymerase chain reaction (RT-PCR) was performed using RNA extracted from MRSA N315 by specific primers to test whether *tlpp1, tlpp2,* and *tlpp3* were co-transcribed (Fig 2A and S5 Table). The comparison of the RT-PCR results with the template of the genomic DNA or RNA only revealed that *tlpp1, tlpp2,* and *tlpp3* were co-transcribed from the *tlpp1* promoter (Fig 2B). We further examined the influence of β-lactams on *tlpp* expression. Reverse-transcription quantitative PCR (RT-qPCR) showed that the mRNA levels of *tlpp1, tlpp2,* and *tlpp3* were upregulated in N315 after treatment with the subinhibitory concentrations of OXA (Fig 2C). Western blot analysis demonstrated that the protein levels of Tlpps in both N315 total cell lysates (Fig 2D and 2E) and the culture supernatant (Fig 2F and 2G) increased in a dose-dependent manner after OXA treatment was administered. Tlpp expression was also upregulated by N315 in a dose-dependent manner after MET treatment was given (S2 Fig). Furthermore, the Tlpp expression by N315 increased in a time-dependent manner after OXA treatment was administered (Fig 2H and 2I). These results verified that MRSA Tlpps could be released from the bacteria and secreted to the culture, and their production was influenced by the subinhibitory concentrations of β-lactam antibiotics.

### β-lactam-induced Tlpps triggered proinflammatory cytokine production by macrophages

In Gram-negative bacteria, the cell wall-associated lipopolysaccharides are the main molecules involved in activating the innate immune system of hosts via TLR4 interaction [23], while in Gram-positive bacteria, the releasable Lpps may be the main factor that exert a similar function by triggering the TLR2-MyD88-NF-κB signaling pathway, thereby inducing proinflammatory cytokines [17,19]. To determine whether the β-lactam-induced MRSA Tlpps involved in the innate immune activation, we cultured mouse RAW264.7 macrophages in media containing 5% culture supernatant of N315 post-treated with different concentrations of OXA for 6 h. We then determined the levels of proinflammatory cytokines, such as IL-6 and TNFα, in the cell-cultured media (DMEN). The results revealed that IL-6 and TNFα levels were gradually increased by macrophages stimulated by the culture supernatant of N315 treated with OXA in a dose-dependent manner (Fig 3A and 3B). Then, we wondered whether the MRSA Tlpps promoted the cytokine expression in macrophages, a *tlpps* deletion mutant (N315Δ*tlpps*) with a pYT3-Δ*tlpps* plasmid and a tlpps-overexpressing strain (N315Δ*tlpps*/pLI-*tlpps*) with a pLI-*tlpps* plasmid (Fig 3C and S3 Fig) were constructed for macrophage infection. Results showed that the production of IL-6 and TNFα in macrophages significantly decreased after treated with N315Δ*tlpps* at a multiplicity of infection (MOI) of 30 compared with those of the wild-type N315 administered. By contrast, higher levels of IL-6 and TNFα were detected in macrophages treated with N315Δ*tlpps*/pLI-*tlpps* but not with N315Δ*tlpps*, which carried an empty pLI50 vector (Fig 3D and 3E). Similar results were observed when the bacterial culture supernatant was used (Fig 3G and 3H) because of the loss of Tlpp expression in the supernatant of N315Δ*tlpps* (Fig 3F), indicating that the increased levels of IL-6 and TNFα by macrophages depended on the expression of N315 Tlpps that responded to β-lactams in a dose-dependent manner. However, the recombinant Tlpp1 purified from *Escherichia coli* (Fig 1C) exhibited no effect on the levels of IL-6 and TNFα by macrophages (S4 Fig), suggesting that a correctly triacylated Tlpp or a long-chain N-acylated Lpp was needed for the recognition by TLR2-TLR1 receptors to trigger immune response by macrophages [24].

**Fig 3.**
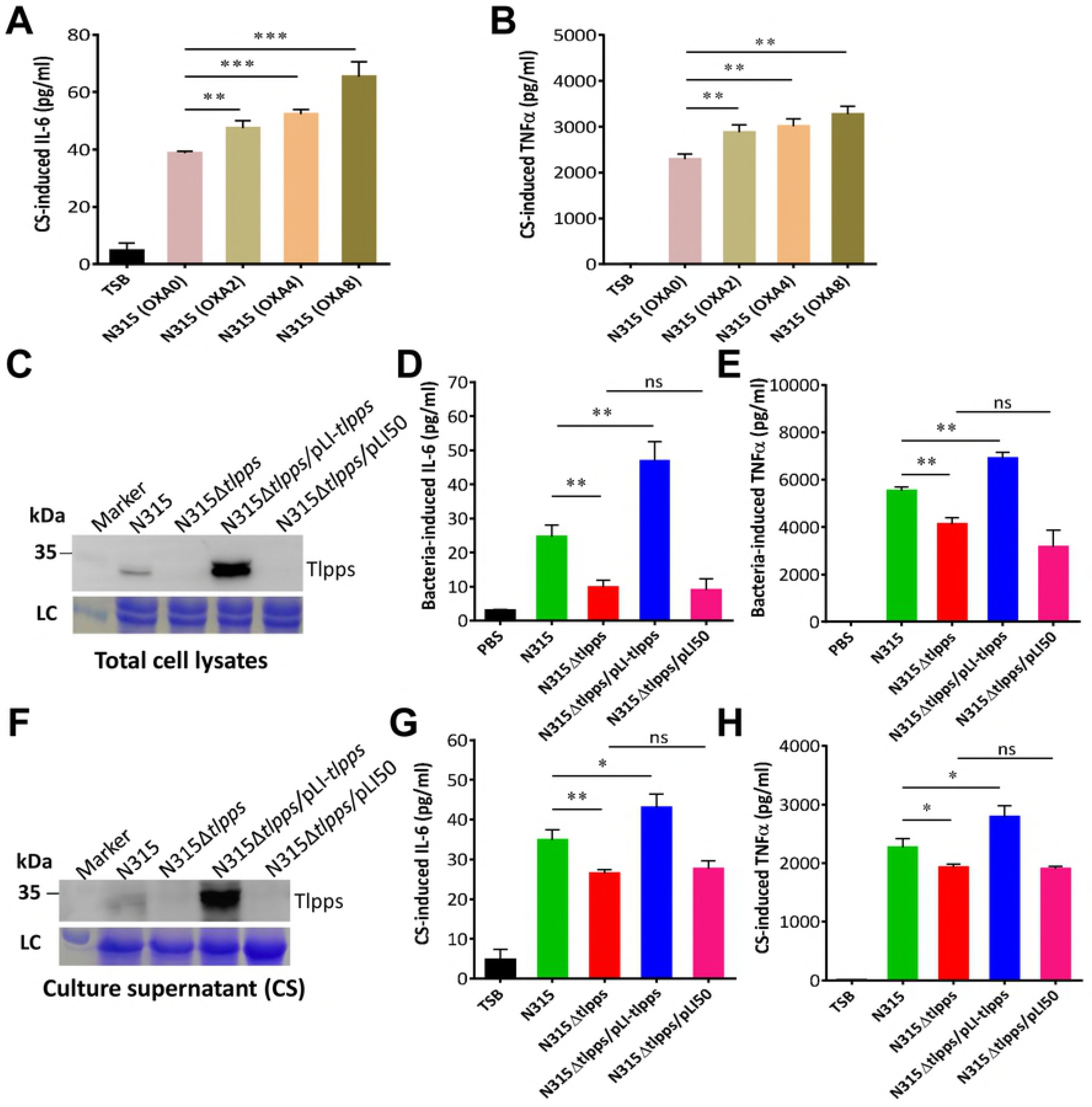
Antibiotic-induced MRSA Tlpps stimulated proinflammatory cytokine production by macrophages. IL-6 (A) and TNFα (B) levels increased by macrophages stimulated by 5% culture supernatant (CS) of N315 treated with different concentrations of OXA as indicated for 6 h. The cell culture of TSB stimulated cells served as the negative control. (C) Western blot analysis of the Tlpp proteins in N315Δ*tlpps* and *N315Δ*tlpps*/pLI-*tlpps*.* Wild type N315 and N315Δ*tlpps*/pLI-50 served as the controls. IL-6 (D) and TNFα (E) levels elevated by macrophages stimulated with N315, N315Δ*tlpps*, N315Δ*tlpp*/pLI-*tlpps*, and N315Δ*tlpps*/pLI50 at the MOI of 30. (F) Western blot analysis of the Tlpp proteins in the culture supernatant of N315Δ*tlpps* and N315Δ*tlpps*/pLI-*tlpps*. IL-6 (G) and TNFα (H) levels increased by macrophages treated with 5% culture supernatant (CS) of N315, N315Δ*tlpps*, N315Δ*tlpp*/pLI-*tlpps*, and N315Δ*tlpps*/pLI50, respectively. The experiments were conducted in triplicate. ns represented no statistical significance, * *P* < 0.05, ** *P* < 0.01, *** *P* < 0.001.

### Tlpps contributed to the virulence of MRSA

Consistent with the observed immune response by macrophages, the levels of IL-6 and TNFα in mice 6 h post-challenge with 1 × 10^7^ N315Δ*tlpps* cells by tail vein injection were significantly lower than those of N315 was administered (Fig 4A and 4B). The complementary overexpression of *tlpps* (N315*Δtlpps*/pLI-*tlpps*) strain stimulated even higher levels of IL-6 and TNFα in mice, but not of the empty pLI50 plasmid-carrying strain (N315Δ*tlpps*/pLI50). We also investigated whether the *tlpp* cluster of MRSA was associated with the bacterial burden in a mouse model. The mice were infected intravenously with pGFP plasmid-transformed N315 and N315*Δtlpps* for 5 days (S6 Table), and bacterial colonization was tracked through an animal imaging system. The fluorescence intensity of the GFP in the murine organs (i.e., heart, lung, liver, spleen, and kidney) was measured, and the results indicated that the radiant efficiency in the kidneys of the mice injected with N315 was significantly higher than those infected with N315Δ*tlpps* (Fig 4C and 4D). Consistent with the radiant efficiency, the bacterial loads in the kidneys of the N315-infected mice were also significantly higher than that of the N315Δ*tlpps*-infected ones (Fig 4E). Overall, these data suggested that the systemic inflammatory response in MRSA infection was associated with Tlpps, and MRSA Tlpps contributed to bacterial colonization and virulence.

**Fig 4.**
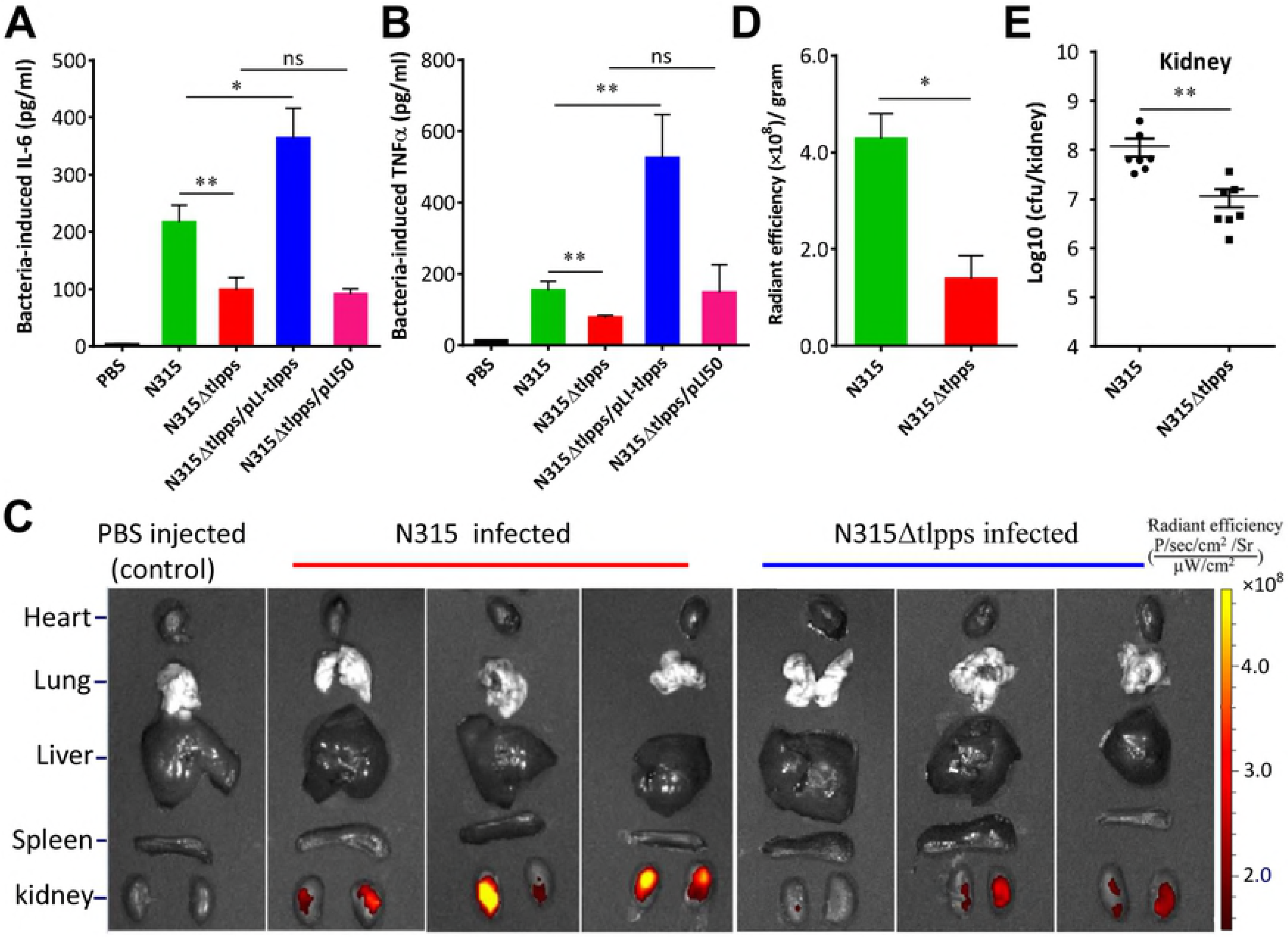
Tlpps contributed to the inflammatory response and bacterial colonization in MRSA infection. IL-6 (A) and TNFα (B) levels in mouse sera determined by ELISA. BALB/c mice were infected by tail vein injection with 1 × 10^7^ of N315, N315Δ*tlpps*, N315Δ*tlpps*/pLI-*tlpps*, and N315Δ*tlpps*/pIL50, respectively. The levels of IL-6 and TNFα in mouse sera were determined 6 h post-infection. Number of mice used, *n* = 3. Phosphate-buffered saline (PBS, pH 7.2) served as the negative control. (C) Organ distribution of N315 and N315Δ*tlpps*. BALB/c mice were infected with 1 × 10^7^ μgFP plasmid transformed N315 and N315Δ*tlpps*, the radiant efficiency of the indicated organs was measured with IVIS^®^ Lumina LT system. (D) Tlpps contributed to the MRSA colonization. The radiant efficiency of N315-infected organs was higher than that of N315Δ*tlpps*-infected ones. (E) Bacterial loads in kidney. BALB/c mice (*n* = 7 for each group) were respectively infected with 1 × 10^7^ of N315 and N315Δ*tlpps* for 5 days, bacteria were recovered and counted from kidneys. ns represented no significance, * *P* < 0.05, ** *P* < 0.01.

### β-lactam-antibiotic-stimulated Tlpps promoted the pathogenesis of MRSA

We determined whether β-lactam-stimulated Tlpps enhanced the pathogenesis of MRSA. A mouse subcutaneous infection model was used to evaluate the contribution of the subinhibitory concentrations of OXA or Tlpps to skin and soft tissue infections. The mice were subcutaneously injected in both flanks with 5 × 10^7^ OXA-treated N315 and N315Δ*tlpps* cells and intraperitoneally injected with 1 μg of OXA per gram weight twice a day for 14 days. The course of infection was monitored every day. The untreated N315- and N315Δ*tlpps*-infected and PBS-injected mice served as the controls. The abscesses caused by the OXA-treated N315 was significantly larger than those caused by the OXA-treated N315Δ*tlpps*, untreated N315, and untreated N315Δ*tlpps* (Fig 5A), and this observation was further shown in the photographs of the skin lesions (Fig 5B and S5 Fig). Histological examinations indicated that the skin of the OXA-treated N315-challenged mice exhibited less extensive inflammation with leukocyte infiltration, destroyed skin structure, and more flake-like abscess-formation than those of the untreated N315-infected and PBS-injected mice (Fig 5C). By contrast, the skin of the OXA-treated N315Δ*tlpps*-challenged mice displayed more leukocyte infiltration and sporadic abscess formation than that of the untreated N315Δ*tlpps*-infected and PBS-injected mice. The <u>corium layer</u> of the N315-challenged and PBS-injected mice showed an extensive inflammation with leukocyte infiltration, although abscess formation was not observed compared with that of the N315Δ*tlpps*-infected and PBS-injected mice. These pathological phenomena might be caused by the β-lactam-stimulated MRSA Tlpps, which stimulated the IL-6 and TNFα levels in mice (Fig 5D and 5E), thereby silencing the immune responses through granulocytic and monocyticmyeloid-derived suppressor cells induced by IL-6 [17], increasing immune cells death due to the tremendous TNFα release [24], and promoting bacterial colonization and abscess formation. Overall, these data confirmed that β-lactam-stimulated Tlpps worsened the MRSA infections.

**Fig 5.**
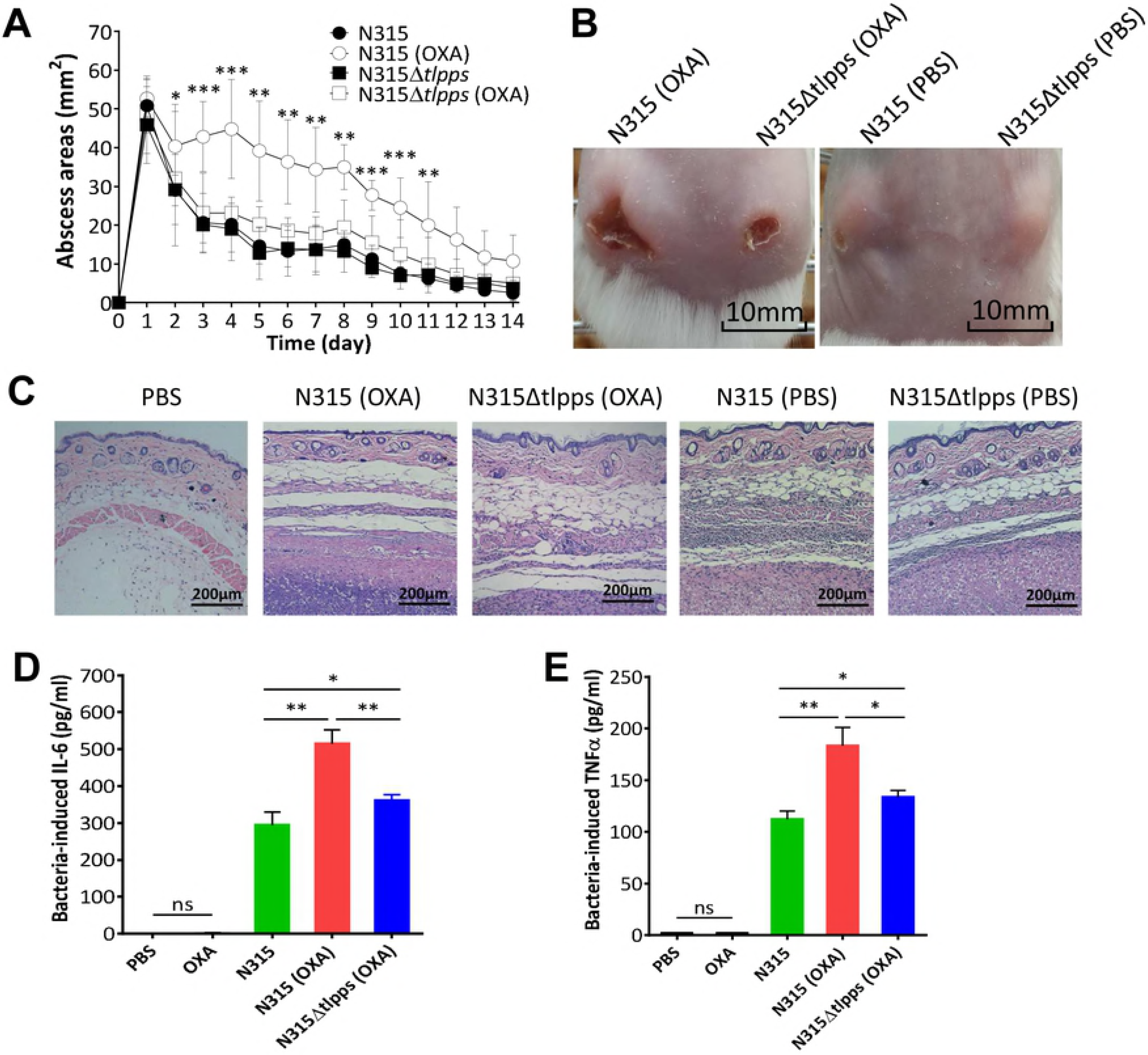
β-lactam treatment enhanced the pathogenesis of MRSA in the mouse subcutaneous infection model. (A) Mice were injected subcutaneously with 5×10^7^ bacterial cells. Abscess areas were measured daily. (B) Representative abscesses 7 days after infection. (C) Representative histological examinations (H&E stain) of the infected mouse skin. IL-6 (D) and TNFα (E) levels in mouse sera determined by ELISA. BALB/c mice were infected by tail vein injection with 1 × 10^7^ of N315, OXA-treated N315, and OXA-treated N315Δ*tlpps*, respectively. The levels of IL-6 and TNFα in mouse sera were determined 6 h post-infection. Number of mice used, *n* = 3. PBS and OXA served as the controls. ns represented no significance, * *P* < 0.05, ** *P* < 0.01, *** *P* < 0.001.

### β-lactam-stimulated Tlpp expression in MRSA was SarA dependent

β-lactams as antibiotics block the cell wall synthesis of bacteria to exert antimicrobial effects. By contrast, β-lactams as inductors may trigger global regulatory networks to modulate virulence in *S. aureus* [13]. RT-qPCR revealed that the expression of global regulators, including *sarA, agrA, RNAIII, rot, ccpA,* and *saeR*, increased in OXA-treated N315 compared with those in the untreated ones (Fig 6A). *SarA* and *agrA* were the most altered regulators, whereas *sigB* was unchanged, which was consistent with the Western blot results of SigB stimulated with different OXA concentrations (S6 Fig). AgrA is a downstream regulator of SarA [25]. In this study, we examined whether SarA was upregulated in N315 upon OXA treatment. Western blot analysis indicated that both SarA and Tlpps increased in a dose-dependent manner in response to the β-lactam antibiotic treatment (Fig 6B, 6C and S2 Fig). The deletion of *sarA* reduced the Tlpps levels in both N315 and its culture supernatant (Fig 6D and S7 Fig). The *sarA* overexpressing strain (N315Δ*sarA*/pLI-*sarA*) produced even more Tlpps compared with the wild type N315, whereas the empty pLI50 plasmid-carrying strain (N315Δ*sarA*/pLI50) did not. Consistent with the decreased Tlpps in N315Δ*sarA*, N315Δ*sarA* or its culture supernatant stimulated less IL-6 and TNFα expression in RAW264.7 cells than N315 or its culture supernatant did. By contrast, N315Δ*sarA*/pLI-*sarA* or its culture supernatant induced more IL-6 and TNFα expression than the wild-type N315 or N315Δ*sarA*/pLI50 did (S8 Fig). The IL-6 and TNFα levels in mice challenged with N315Δ*sarA* decreased compared with those in the mice challenged with N315. N315Δ*sarA*/pLI-*sarA* stimulated even higher cytokine levels in mice compared with those of N315 or N315Δ*sarA* induced (Fig 6E and 6F). Taken together, these data indicated that the β-lactam-stimulated Tlpp expression in MRSA was SarA dependent.

**Fig 6.**
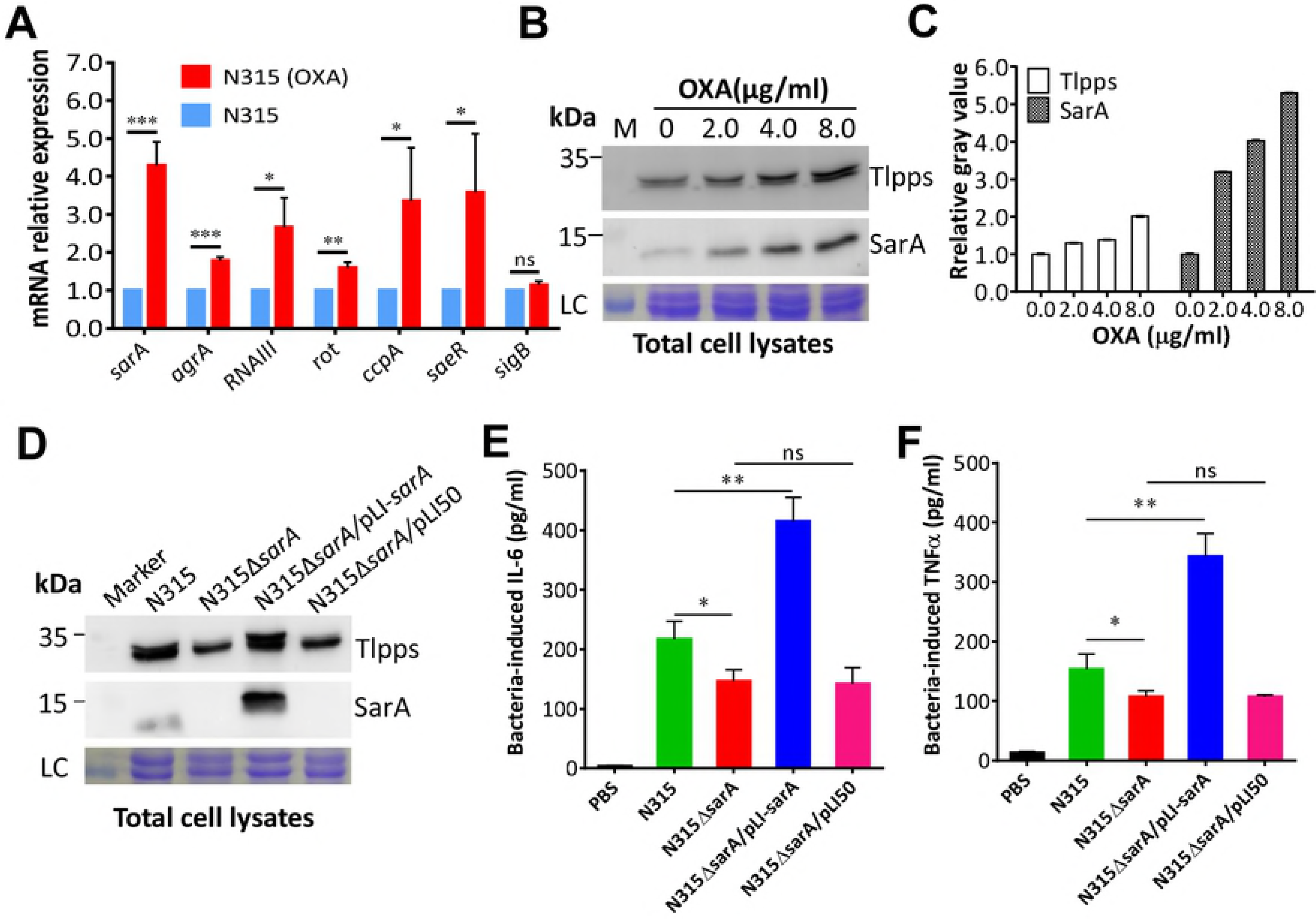
SarA affected the Tlpp expression in MRSA. (A) RT-qPCR detection of the expression levels of global regulators in MRSA with or without OXA treatment. The expression of each gene in N315 was normalized to that of the 16S *rRNA* gene and adjusted to 1.0, the relative expressions of regulators in OXA-treated N315 were indicated. The experiments were repeated for three times. (B) Western blot analysis of SarA and Tlpp proteins in N315 treated with different concentrations of OXA. (C) Evaluation of gray value of Tlpp and SarA in N315 after OXA treatment. (D) The deletion of *sarA* decreased the Tlpp expression in the N315. SarA mutant stimulated less IL-6 (E) and TNFα (F) production in mice. Mice were infected with 1×10^7^ of N315, N315Δ*sarA*, N315Δ*sarA* /pLI-*sarA*, and N315Δ*sarA*/pLI50. The levels of IL-6 and TNFα in mouse sera were determined by ELISA 6 h post-infection. The experiments in duplicate were conducted for three times. ns represented no statistical significance, * *P* < 0.05, ** *P* < 0.01, *** *P* < 0.001.

### SarA directly controlled the β-lactam-stimulated Tlpp expression in MRSA

To investigate the effect of SarA on β-lactam-stimulated Tlpp expression, we constructed a reporter vector (pOS1-*tlpps*^P^) containing the *tlpp* promoter-controlled *lacZ* gene (S6 Table) and performed β-galactosidase assay by transforming pOS1-*tlpps*^P^ into the MRSA strains N315 and N315Δ*sarA*. The results revealed that the β-galactosidase activity was significantly lower in the *sarA* mutant than that in N315. Moreover, the β-galactosidase activity presented no significant change in the *sarA* mutant after OXA treatment compared with the untreated N315Δ*sarA* (Fig 7A). However, OXA treatment significantly increased the β-galactosidase activity in N315, and this increase was consistent with the increasing Tlpp in MRSA strains (Fig 7A, Fig 1A and 1B). Consistent with the β-galactosidase assay results, Western blot analysis demonstrated that the Tlpp expression in the *sarA* mutant could not respond to β-lactam simulation (Fig 7B and 7C), suggesting that SarA controlled the Tlpp expression in response to the β-lactam antibiotic induction.

**Fig 7.**
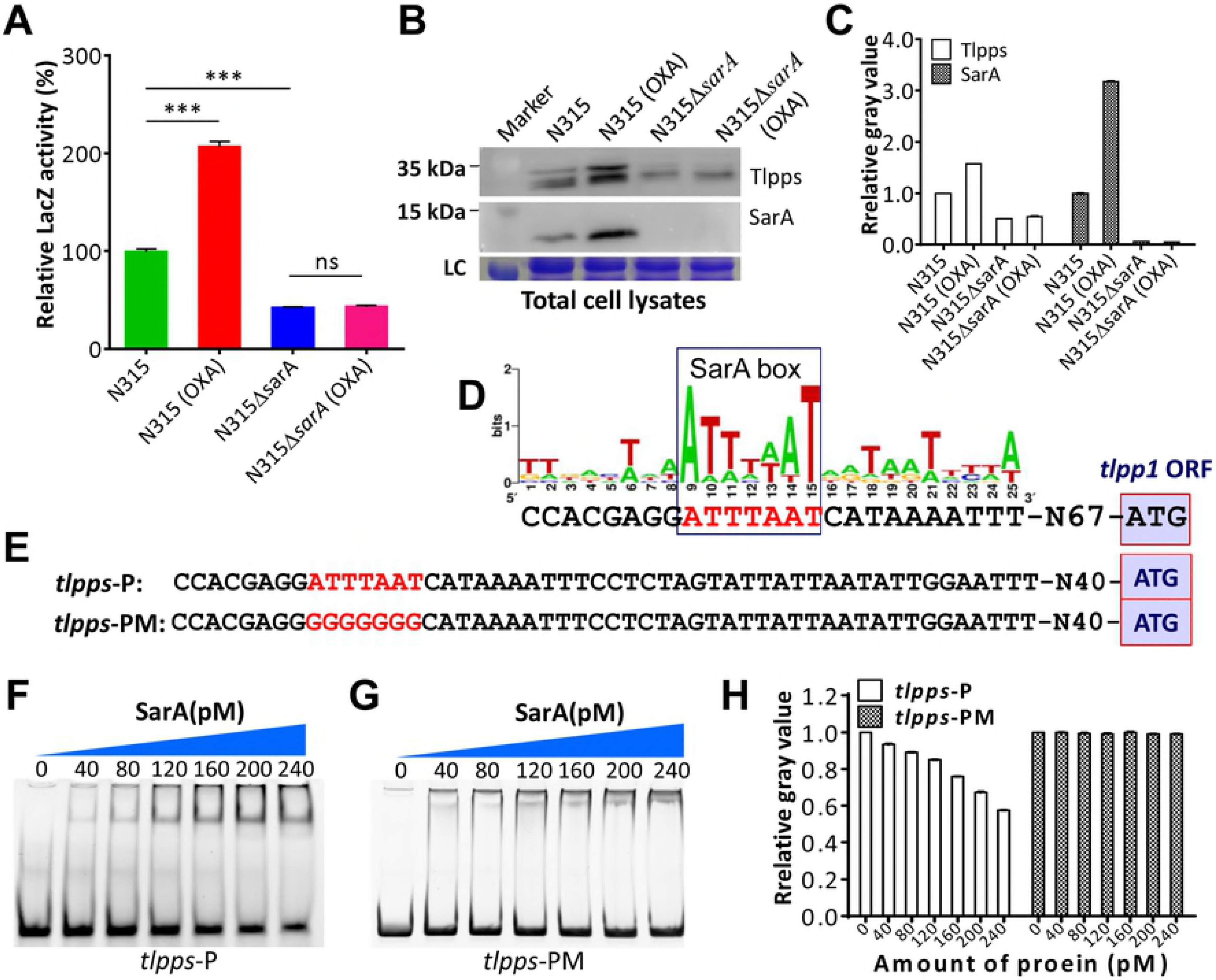
SarA bound to the *tlpp* cluster promoter region to control the β-lactam-stimulated Tlpp expression in MRSA. (A) β-galactosidase assay. The pOS1-*tlpps*^P^ reporter plasmid was transformed into N315 and N315Δ*sarA*, respectively. The LacZ activity was detected and represented as mean ± S.D. (*n* = 3). ns indicated no significance, \*\*\**P* < 0.001. (B) SarA controlled β-lactam-stimulated Tlpp expression in N315. (C) Evaluation of gray value of Tlpps and SarA in N315 and N315Δ*sarA* with or without OXA treatment. (D) Predicted SarA box in the promoter regions of the *tlpp* cluster. (E) Mutation of AT-rich in the *tlpp* SarA box for EMSA experiment. (F) EMSA. Interaction between wild-type *tlpp* promoter region (*tlpps*-P) and SarA proteins was detected. (G) EMSA with *tlpp* promoter mutant (*tlpps*-PM). (H) Evaluation of gray value of the free probe in each lane using ImageJ software. The value of free probe in the first lane (0 μg protein) was adjusted to 1.0, and the relative gray values in other lanes were calculated and indicated.

A global regulator can recognize specific motifs in the promoter regions of a certain gene, thereby controlling the gene expression [13,26,27]. We analyzed the binding motif of SarA [28,29] in the promoter regions of *tlpps* and found a typically predicted SarA box (Fig 7D). The recombinant His-tagged SarA was prepared and purified from *E. coli* (S7C Fig), and the electrophoretic mobility shift assay (EMSA) showed that the recombinant SarA proteins bound to the *tlpp* cluster promoter region that carried the putative SarA binding box (Fig 7E, 7F and 7H). No shifting band was observed when the AT-rich SarA box was mutated to become GC-rich (Fig 7E, 7G and 7H). These data indicated that *S. aureus* SarA could directly bind to the *tlpp* cluster promoter region, thereby upregulating the *tlpp* expression in the presence of β-lactams. The AT-rich motif (ATTTAAT) in the promoter regions of *tlpps* is essential for SarA binding and regulating.

## Discussion

MRSA is distinct from MSSA in terms of the acquisition of a genetic element called staphylococcal cassette chromosome *mec* (SCC*mec*) in which *mecA* encodes an alternative penicillin-binding protein 2a (PBP2a) with a low affinity for β-lactams [21]. As such, MRSA strains are resistant to nearly all β-lactam antibiotics [6]. As antibiotics, β-lactams bind to penicillin-binding proteins (PBPs) and inhibit the transpeptidation and transglycosylation of the cell wall, resulting in a weakened cell wall and inducing cell lysis and death [30]. This type of antibiotics, particularly cephalosporins and β-lactam/β-lactamase inhibitor combinations, has been empirically used for clinical treatments of infectious diseases [31]. The subinhibitory concentrations of antistaphylococcal agents might arise because of antibiotic-resistant microorganisms or pharmacokinetics of antibiotics [13,31]. For MRSA infections, which are not initially recognized, β-lactams are not only ineffective in the treatment of infections but also likely contributing to poor outcomes by enhancing the pathogenicity of MRSA. Nonetheless, the underlying mechanisms remain obscure [9]. In this study, we showed that a three-gene constituent *tlpp* cluster upregulated in response to β-lactam induction in a dose- and time-dependent manner. This *tlpp* cluster, belonging to a DUF576 protein family of unknown function [19,22], was widely distributed among the major prevalent MRSA clones (Fig 1B and S4 Table). Tlpps could be upregulated after treatment with nearly all β-lactam antibiotics, such as penicillin (oxacillin and methicillin), cephalosporins (cefoxitin), and carbapenems (imipenem and meropenem), but not with vancomycin, kanamycin, and erythromycin (Fig 1A).

In addition to antimicrobial activity, signal induction may be implemented by the subinhibitory concentrations of β-lactams. They actively promote *S. aureus* biofilm formation [3], induce PBP2a to reduce peptidoglycan crosslinking in MRSA [6], and enhance virulence factors, such as alpha-toxins, PVL, SPA, and enterotoxins [2,32–34]. In contrast to β-lactam-stimulated SPA and PVL, which have a controversial pathogenic role in *S. aureus* [2], the Lpps of *S. aureus* are crucial players in alerting the host immune system by recognizing TLR2/TLR1 or TLR2/TLR6 receptors [24,35]. In general, the Lpps of Gram-positive bacteria are anchored in the outer leaflet of the cytoplasmic membrane [17]. We observed that the β-lactam-stimulated MRSA Tlpps could be released; Tlpps could be detected in both cell lysates and the culture supernatant through Western blot analysis (Fig 2D, 2F, 3C and 3F). MRSA Tlpps could induce the production of IL-6 and TNFα proinflammatory cytokines of macrophages which stimulated with the culture supernatant of N315 post-treated with different OXA concentrations (Fig 3A and 3B). Although the *tlpp* deletion mutant (N315Δ*tlpps*) infection induced less IL-6 and TNFα production in mice compared with the wild-type N315 was administrated, OXA-treated N315Δ*tlpps*-infected mice still produced higher IL-6 and TNFα cytokines compared with those of the untreated N315-challenged mice (Fig 5D and 5E), suggesting that other mechanisms might be involved in the immune system modulation by β-lactam-treated MRSA. For instance, β-lactam-promoted PBP2a induction can diminish the peptidoglycan crosslinking, thereby enhancing phagocytic degradation and detection and resulting in strong IL-1β production [6].

In addition to stimulating the immune system, increasing the pathogenicity of MRSA was attributed to β-lactam-stimulated Tlpps. In comparison with wild-type N315, the N315Δ*tlpps* strain significantly decreased the bacterial burden in mouse kidneys (Fig 4C, 4D and 4E). β-lactam antibiotic treatment exacerbated MRSA infections in the mouse skin infection model, and the histological examinations of the OXA-treated N315-challenged mouse skin displayed less extensive infiltration with leukocytes, destroyed skin structure, and easily promoted abscess formation (Fig 5). A possible explanation is that the induced IL-6 and TNFα expression by β-lactam-stimulated MRSA Tlpps silenced the innate immune responses [24], thereby facilitating MRSA colonization and infection. Our findings might indicate that MRSA infections were associated with higher morbidity and mortality than MSSA [7,36].

β-lactams can induce PVL expression in *S. aureus* by interfering with PBP1 and triggering SarA and Rot global regulators [2]. Our results showed that SarA and AgrA were the most upregulated regulators in MRSA N315 after OXA treatment (Fig 6A). The deletion of SarA (N315Δ*sarA*) failed to upregulate *tlpps* even under OXA treatment (Fig 7A and 7B), indicating that the β-lactam-induced Tlpp expression in MRSA was SarA controlled. EMSA data revealed that the regulation of SarA on Tlpp expression was direct (Fig 7F). However, further investigations should be performed to clarify how β-lactams trigger the SarA expression.

In conclusion, this work focused on the function and regulation of a new *tlpp* cluster in response to the induction of subinhibitory concentrations of β-lactams. We demonstrated that the increased Tlpps in MRSA significantly enhanced inflammatory response by triggering IL-6 and TNFα levels in vitro and in vivo, thereby possibly contributing to bacterial pathogenicity and worsening the outcome of MRSA infection by reducing host immune responses and promoting bacterial colonization. β-lactam-stimulated MRSA Tlpps was SarA dependent, and the upregulation in Tlpp expression after β-lactam treatment was directly controlled by the global regulator SarA (Fig 8). Nonetheless the pathways leading to the SarA expression trigged by β-lactam antibiotics remain unknown. Our data support the recommendation to clinicians that the discreet usage of β-lactams which possibly worsened the clinical outcome of MRSA infections.

**Fig 8.**
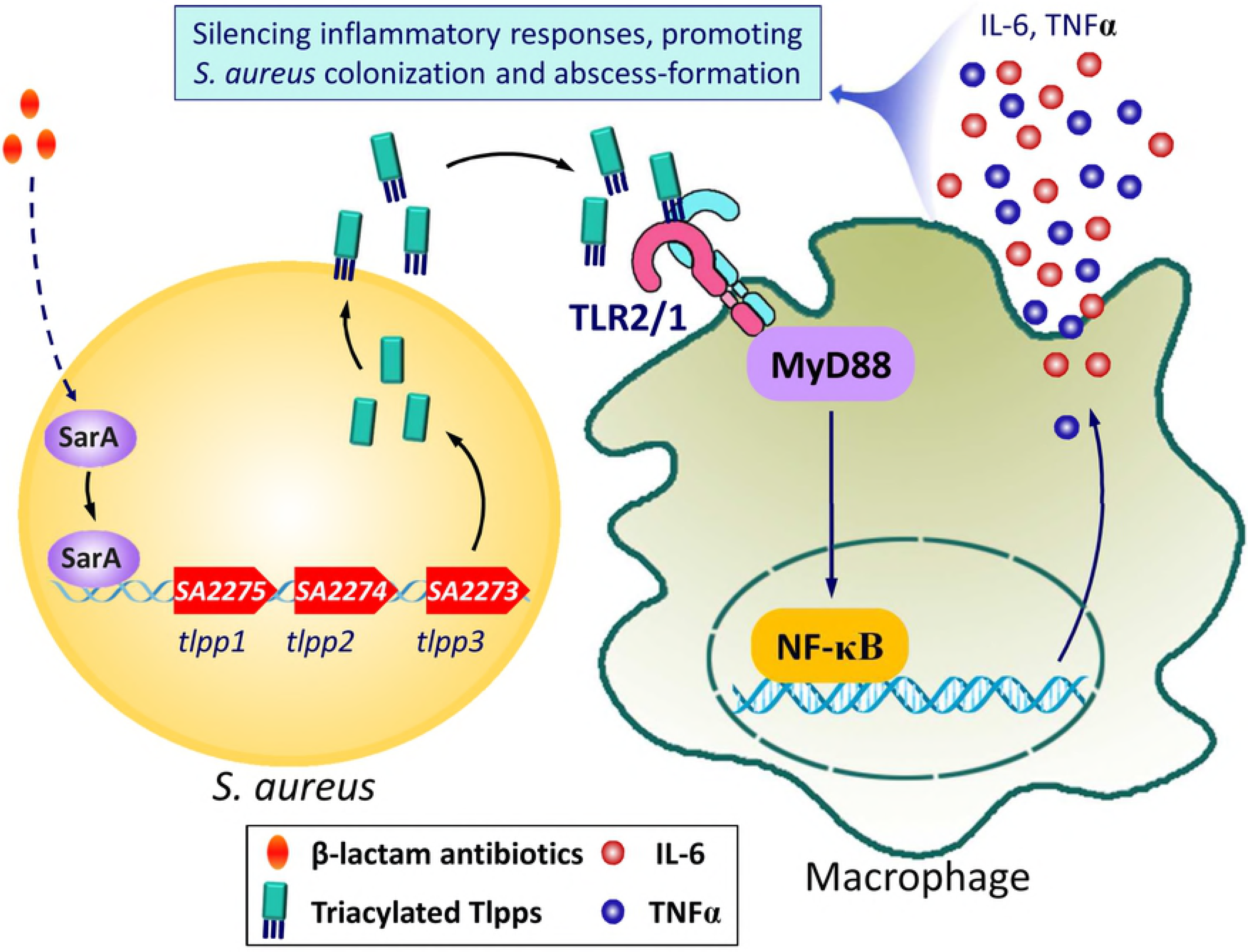
Schematic of the Tlpp function and its upregulation in MRSA after β-lactam treatment.

## Materials and methods

### Ethics statement

BALB/c mice were purchased from Laboratory Animal Center of Army Medical University (Third Military Medical University). All animal experiments were approved by the Army Medical University Institutional Animal Care and Use Committee (protocol #SYXK-PLA-20120031). All animal experimental procedures were performed in accordance with the Regulations for the Administration of Affairs Concerning Experimental Animals approved by the State Council of People`s Republic of China.

### Bacterial strains and plasmids

Bacterial strains and plasmids used in this study were listed in S6 Table. *E. coli* strains DH5α and BL21 (DE3) were cultivated in Luria Broth (LB) medium (Oxoid). *S. aureus* strains grown in Brain Heart Infusion (BHI) or Tryptic Soy Broth (TSB) medium (Oxoid).

### Antibiotic susceptibility tests

Antibiotic susceptibility was determined using broth microdilution methods according to the protocols recommended by the Clinical and Laboratory Standards Institute (CLSI, 2017) [37]. The antibiotic susceptibility results for all strains were listed in S2 Table.

### Preparation of recombinant Tlpp1 and SarA

Recombinant Tlpp1 and SarA proteins were prepared in our laboratory. In brief, the gene encoding for Tlpp1 or SarA was amplified from the genomic DNA of N315 by PCR, and cloned into pET28a(+) vector to construct the pET28a-*tlpp1* and pET28a-*sarA* plasmids. Then, the recombinant plasmids were transformed into *E. coli* BL21(DE3) for the expression of Tlpp1-6×His and SarA-6×His fusion proteins. Cells were grown in LB with 100 μg/ml of ampicillin at 37 °C. Isopropyl-D-thiogalactopyranoside (IPTG, 5 μM) was added once an OD600 of 0.6 was achieved. The cells were cultured at 22 °C for another 6 h. The cells were then centrifuged at 10,000 × g for 15min, washed once with PBS, resuspended in the lysis buffer (50 mM Tris·Cl, 0.15 M NaCl, 1 mM phenylmethanesulfonyl fluoride (PMSF), 0.5mg/ml lysozyme, pH 8.0) and lysed using ultrasonic method. The recombinant proteins in the supernatant were purified by Ni-NTA affinity chromatography (GE Healthcare) and identified by Western blot.

### Preparation of protein-specific polyclonal antibodies

Female BALB/c mice (6-8 weeks) were immunized subcutaneously with 40 μg of recombinant proteins (Tlpp1 or SarA) emulsified with complete Freund’s adjuvant (Sigma, USA) for the first time, and boosted with recombinant proteins (Tlpp1 or SarA) in incomplete Freund’s adjuvant on day 14, 28, and 35, respectively. Seven days after the last immunization, blood samples were collected, the titer of antibodies against Tlpp1 or SarA was determined by ELISA.

### Preparation of the total bacteria proteins and the culture supernatant proteins

Overnight MRSA culture was diluted 1:100 in BHI medium with or without the addition of β-lactam antibiotics, cultivated at 37 °C to an optical density (OD) at 600 nm of 2.0. Then, bacterial cells in 3 ml culture were harvested by centrifugation at 10,000 × g at 4 °C. The cell pellets were washed twice with PBS, resuspended in 1 ml of cold PBS supplemented with 1% β-mercaptoethanol (Sigma, USA) and 1 mM PMSF (Beyotime, China) on ice. Cells were broken by the addition of 0.1-mm diameter zirconia/silica beads, shaking on the Minibeadbeater 16 instrument (Biospec, USA). Cell debris was removed after centrifugation at 10,000 × g for 10 min at 4 °C, and the total cell proteins in 1 ml of the supernatant were precipitated with 7.5% trichloroacetic acid (TCA)/0.2 % deoxycholic acid solution.

The culture supernatant of MRSA stimulated with or without antibiotics was harvested by centrifugation at 10,000 × g for 10 min at 4 °C. The proteins in 1 ml culture supernatant were prepared by TCA precipitation, collected by centrifugation at 15,000 × *g* for 10 min, washed once with ice-cold acetone, dissolved in 60 μl of PBS for use. The protein concentration was determined using the Bradford Protein Assay Kit (Beyotime, China).

### Proteomic analysis

LC-MS/MS was performed to identify proteins induced by β-lactam antibiotics as previously described [38]. MRSA strain N315 was cultured in TSB with or without 2 βg/ml OXA. The total bacteria proteins were separated through SDS-PAGE, and the antibiotic-induced protein band was excised and analyzed through LC-MS/MS by using UltiMate3000 RSLCnano liquid chromatography/Bruker maxis 4G Q-TOF. The resulting peptide mass fingerprints were compared against the ORFs of N315 by using Mascot and Mascot Daemon software (Matrix Science).

### RT-PCR and RT-qPCR

Total MRSA N315 RNA was extracted as previously described [39]. Overnight N315 cultures were 1:100 diluted in TSB containing 2 μg/ml OXA, cultivated at 37 °C to the early exponential phase (OD600 = 1.0). Total RNA was isolated using the TriPure isolation reagent (Roche Applied Science, Germany) after the collected cells were firstly lysed using lysostaphin (Sigma, USA). cDNA was synthesised from 500 ng of total RNA using gene-specific primers and a RevertAid First Strand cDNA Synthesis Kit (Thermo, USA). RT-PCR was used to test whether *tlpp1, tlpp2,* and *tlpp3* were cotranscribed. RT-qPCR was performed to detect the expression levels of the *tlpp* genes (*tlpp1*, *tlpp2* and *tlpp3*) and the global regulators (*sarA, agrA, RNAIII, rot, ccpA, saeR, sigB*) using SsoAdvanced™ Universal SYBR® Green Supermix (Bio-Rad, USA). The relative expression level of all tested genes was normalized to that of the 16S *rRNA*. All primers used are listed in S5 Table.

### Construction of gene mutant strains

The *tlpp* cluster marker-less deletion mutant was constructed using homologous recombinant strategy described previously [39]. Briefly, the *tlpp* cluster deletion plasmid pYT3-Δ*tlpps* (S3A Fig) was constructed by amplifying the upstream and downstream regions of *tlpp* cluster with primer pairs, up-*tlpps* fwd/up-*tlpps* rev and down-*tlpps* fwd/down-*tlpps* rev, and subcloning these fragments into *E. coli-S. aureus* temperature-sensitive shuttle vector pYT3. The recombinant plasmid was identified by DNA sequencing and subsequently transformed into RN4220 by electroporation, then transformed into N315. The resulting N315Δ*tlpps* strain was generated after inducing the integration of plasmid into chromosome at 42 °C and following inducing the plasmid losing at 25 °C. The deletion of *tlpp* cluster was confirmed by PCR and DNA sequencing.

To construct the *tlpps* complementary strain, the *tlpp* cluster containing its potential promoter region was amplified by pLI-*tlpps* fwd/pLI-*tlpps* rev primer pairs, cloned into the expression plasmid pLI50 [40]. Then, the correctly constructed plasmid pLI-*tlpps* was electroporated into RN4220 and then N315Δ*tlpps* to generate N315Δ*tlpp*/pLI-*tlpps* strain. Similar strategy was used to construct N315Δ*sarA*/pLI-*sarA*. The empty pLI50 plasmid transformed N315Δ*tlpps* and N315Δ*sarA* strains served the controls. All primers used are listed in S5 Table.

### Cytokine determination

RAW264.7 (TIB-71™, ATCC) cells were cultured in high glucose DMEM (Thermo Fisher Scientific, USA) added 10 *%* (v/v) fetal bovine serum (Thermo Fisher Scientific, USA) at 37 °C with 5 % CO_2_. For stimulation experiment, RAW264.7 cells (10^6^/well) were either infected with MRSA strain (MOI of 30), or cultured in DMEM medium containing 5% (v/v) bacterial culture supernatant, in a 24-well microtiter plate for 6 h as described [41]. Then, the supernatant was collected, and the levels of IL-6 and TNFα were measured with an ELISA kit following the manufacturer’s instructions (R&D Systems, USA).

To detect the ability of Tlpp proteins in inducing cytokine secretion in vivo, female BALB/c mice were infected via tail vein injection with 1 × 10^7^ CFU of N315, N315Δ*tlpps*, N315Δ*tlpps*/ *pLI-*tlpps*,* N315Δ*tlpps*/pLI50, N315Δ*sarA*, N315Δ*sarA*/pLI-*sarA*, and N315Δ*sarA*/pLI50, respectively. Blood samples were collected 6 h post infection, and the IL-6 and TNFα levels in mouse sera were determined by using ELISA.

### Animal experiments

BALB/c mice were randomly divided into two groups and infected via tail vein injection with 1 × 10^7^ CFU of the GFP-expression plasmid (pGFP) transformed N315 or N315Δ*tlpps*, sacrificed 5 days after infection. Mouse organs (i.e., heart, lung, liver, spleen, and kidney) were isolated and subjected to the determination of GFP fluorescence efficiency in the organs with IVIS^®^ Lumina LT system and analyzed by Living Image 4.4 Software. The bacterial loads in the infected kidneys were also counted via plate dilution assay as described [42].

For skin abscess formation, BALB/c mice were fully anesthetized with 1 % pentobarbital sodium (50 mg/kg) and the back hair was depilated completely with 6 % sodium sulfide (w/v). Then, mice were subcutaneously inoculated with 5×10^7^ CFU of MRSA N315 and N315Δ*tlpps* in both flanks of the murine back as described [43], and then randomly divided into two groups. The mice of the treatment group were intraperitoneally injected with 1 μg of OXA per gram weight twice a day for 14 days. The PBS-injected mice served as the controls. The abscess area assessed by the maximal length × width of the developing ulcer was measured daily.

For histopathological examination, the skin lesions were fixed by 4 % paraformaldehyde, paraffin embed, and stained with hematoxylin & eosin (H&E).

### β-galactosidase assay

A *tlpp* promoter-LacZ reporter plasmid (pOS1-*tlpps*^P^) was constructed by inserting the *tlpp* promoter region into pOS1 vector, and transformed into N315 and N315Δ*sarA*, respectively. Then, pOS1-*tlpps*^P^-carried N315 and N315Δ*sarA* were cultured overnight, diluted 1:100 in BHI, and cultivated at 37 °C to an OD600 of 0.6. Bacterial cells in 200 μl culture were collected by centrifugation, suspended in 100 μl of AB buffer (100 mM KH_2_PO_4_, 100 mM NaCl, pH 7.0), and treated with lysostaphin (20 μg/ml, Sigma) for 15 min at 37 °C. Then, the suspension was added with another 900 μl of ABT solution (AB buffer containing 0.1 % TritonX-100) [44], 50 μl of the solution was mixed with 10 μl of MUG (4-methylumbelliferyl-β-D-galactoside, 4 mg/mL, Sigma) and incubated for 1 h at room temperature. Then, 20 μL of the sample was mixed with 180 μl of ABT solution, and the reaction was monitored at 445 nm with an excitation wavelength of 365 nm. All samples were tested in triplicate. The LacZ activity was normalized to the cell density of OD600, and the relative activity was calculated by setting the LacZ activity from the N315 to 100%. The assay was repeated at least three times.

### Electrophoretic mobility shift assays (EMSA)

The predicted *tlpp* cluster promoter, an AT-rich motif fragment (56 bp), was synthesized using the primer pairs (EMSA-*tlpps*^P^ fwd/EMSA-*tlpps*^PM^ rev) as described [45]. The corresponding mutated GC-rich motif fragment was also synthesized by primer pairs EMSA-*tlpps*^PM^ fwd/EMSA-*tlpps*^PM^ rev and served as the control. Ten picomole of DNA fragment was incubated with a variable amount of recombinant SarA (0 to 240 pM) in a 20 μl reaction mixture containing 10 mM HEPES (pH 7.6), 1 mM EDTA, 2 mM dithiothreitol, 50 mM KCl, 0.05 % Triton X-100, and 5 % glycerol. Binding reactions were equilibrated for 20 min at room temperature before electrophoresis. Reaction mixtures were separated on 6 % native polyacrylamide gel electrophoresis in 0.5 × TBE (Tris/boric acid/EDTA) buffer at 90 V for 2 h at 4 °C. Gels were stained by GelRed dye (Biotium, USA) and observed under UV light. The primers used are listed in S5 Table.

### Statistical analysis

Statistical analysis was carried out using GraphPad Prism 6.0. Unpaired two-tailed student’s t-test, analysis of variance (ANOVA) and Mann-Whitney test were used appropriately to compare the difference between groups. Each experiment was carried out at least three times. Results were presented as mean ± standard deviations (S.D.), and a *P* value less than 0.05 was considered statistically significant. * *P* < 0.05, ** *P <* 0.01, *** *P* < 0.001, and ns represented no significance.

## Supporting Information

**S1 Fig.**
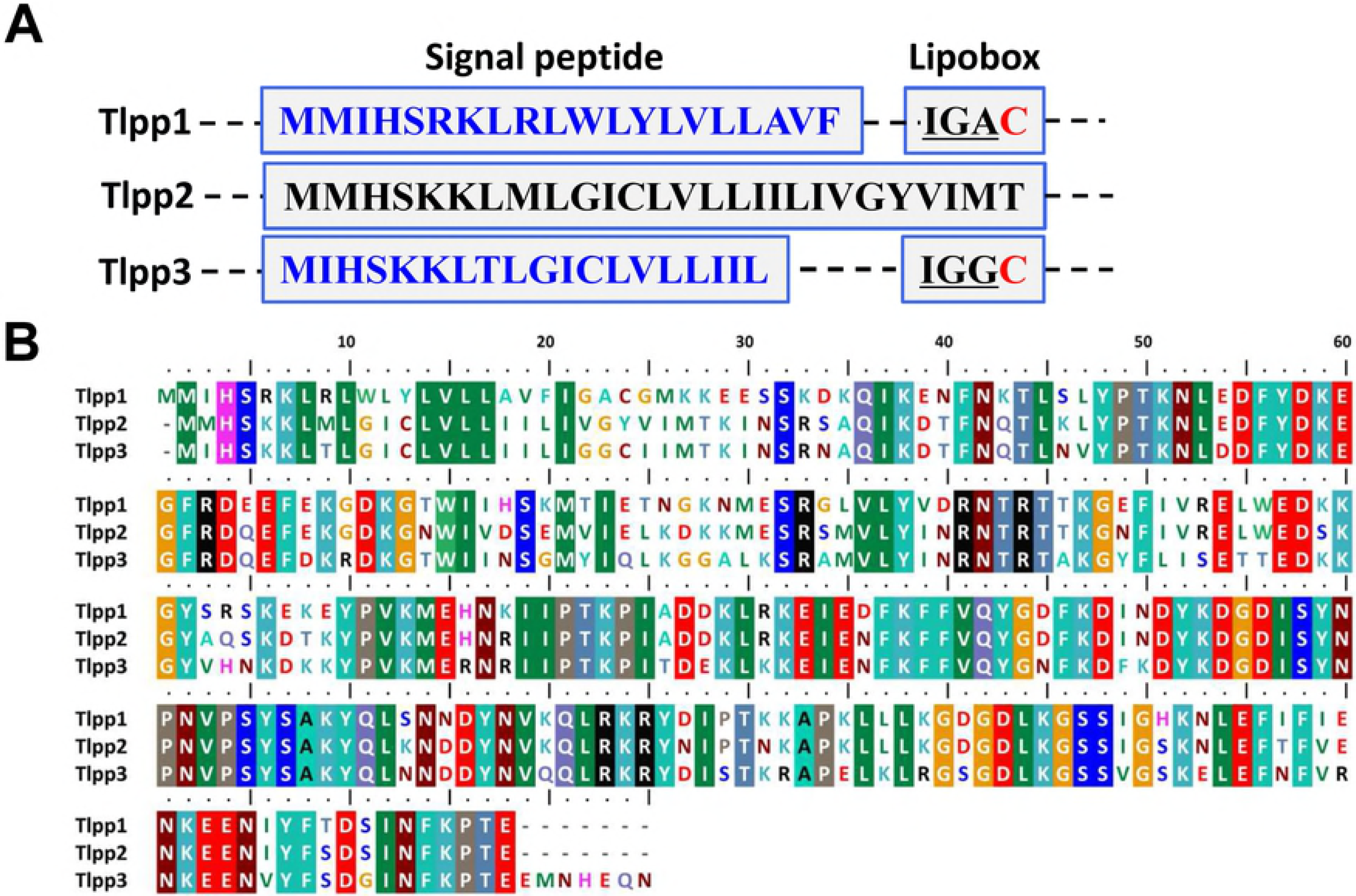
Alignment of the tandem lipoproteins (Tlpps). (A) Alignment of the Tlpp1, Tlpp2, and Tlpp3 signal peptide and the lipobox sequence. (B) Alignment of the Tlpp1, Tlpp2, and Tlpp3 proteins. The amino acid sequences of the indicated protein were retrieved from UniProt (http://www.uniprot.org/), and the alignment was conducted through the BioEdit program. The identical amino acids were colored.

**S2 Fig.**
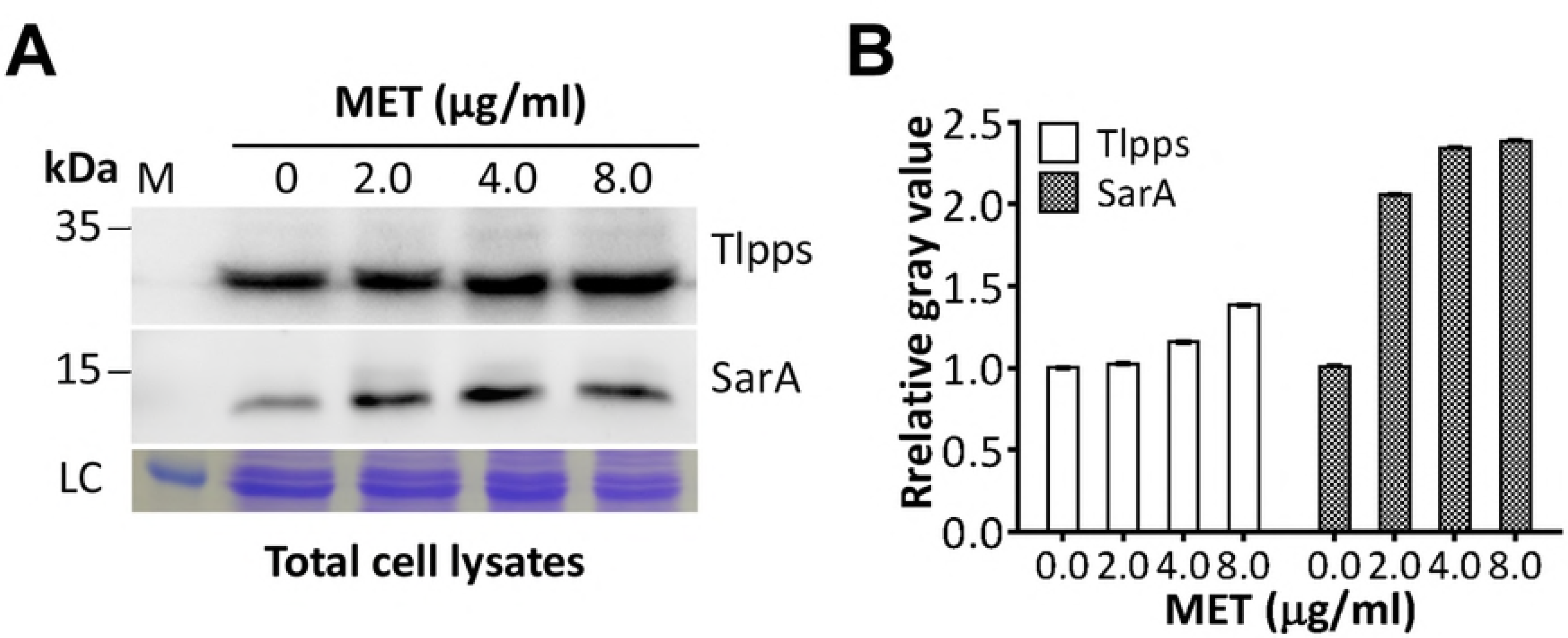
Western blot analysis of Tlpps and SarA in N315 with different concentrations of MET treatment. (A) Tlpps and SarA upregulated in N315 with MET treatment in a dose-dependent manner. The molecular weights of the protein marker (M) were indicated on the left. LC, loading control. (B) Evaluation of gray value of the N315 Tlpps and SarA in each lane using ImageJ software. The relative value of Tlpps or SarA/loading control (LC) in the first lane (0 μg/ml MET) was adjusted to 1.0, and the relative gray values in other lanes were calculated and indicated.

**S3 Fig.**
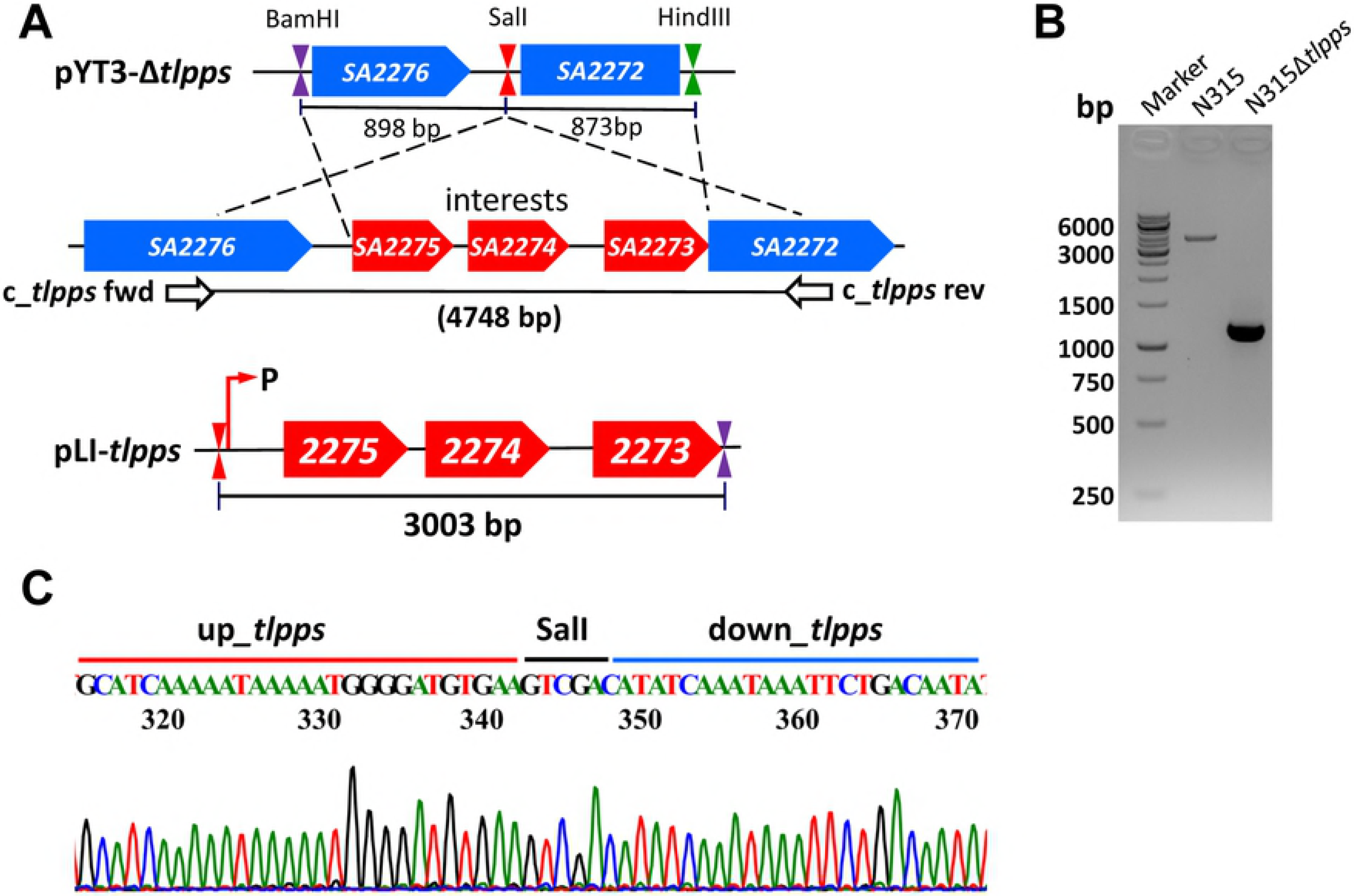
Construction of N315Δ*tlpps* and N315Δ*tlpps*/pLI-*tlpps* strains. (A) Schematics for construction of the *tlpp* marker-less deletion plasmid pYT3-Δ*tlpps* and the complementation plasmid pLI-*tlpps*. (B) Identification of N315Δ*tlpps* by PCR using primer pairs *c_tlpps fwd*/*c_tlpps rev* (S5 Table). (C) Identification of N315Δ*tlpps* by DNA sequencing.

**S4 Fig.**
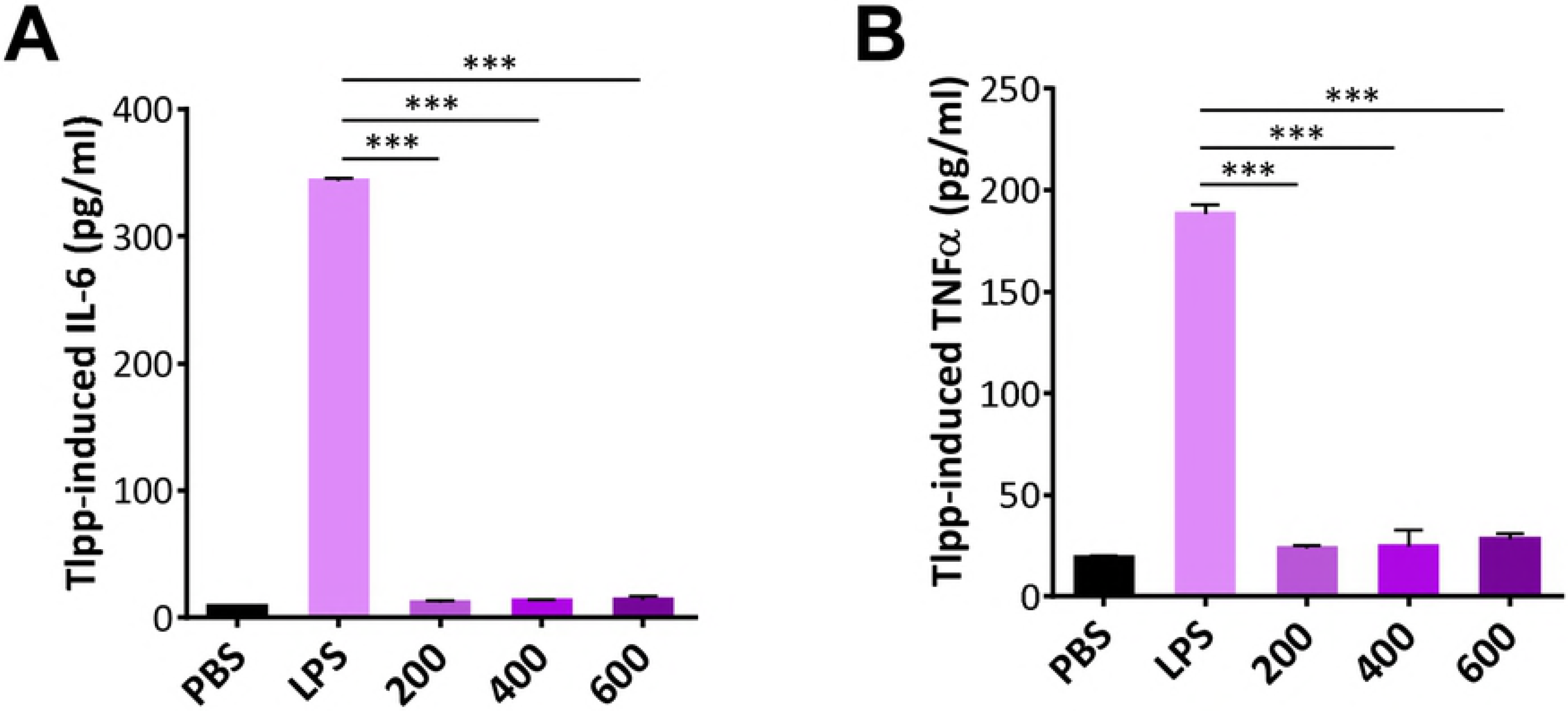
The recombinant Tlpp1 proteins presented no effect on IL-6 and TNFα expression by macrophages. (A) IL-6 and (B) TNFα levels produced by macrophages post-treated with the recombinant Tlpp1 proteins of 200 ng, 400 ng, or 600 μg were given. The cytokine levels were determined by ELISA after 6 h post-treated. LPS (200 ng) administrated was served as the positive control, and PBS-treated served as the negative control. The experiments were conducted in thrice. *** *P* < 0.001.

**S5 Fig.**
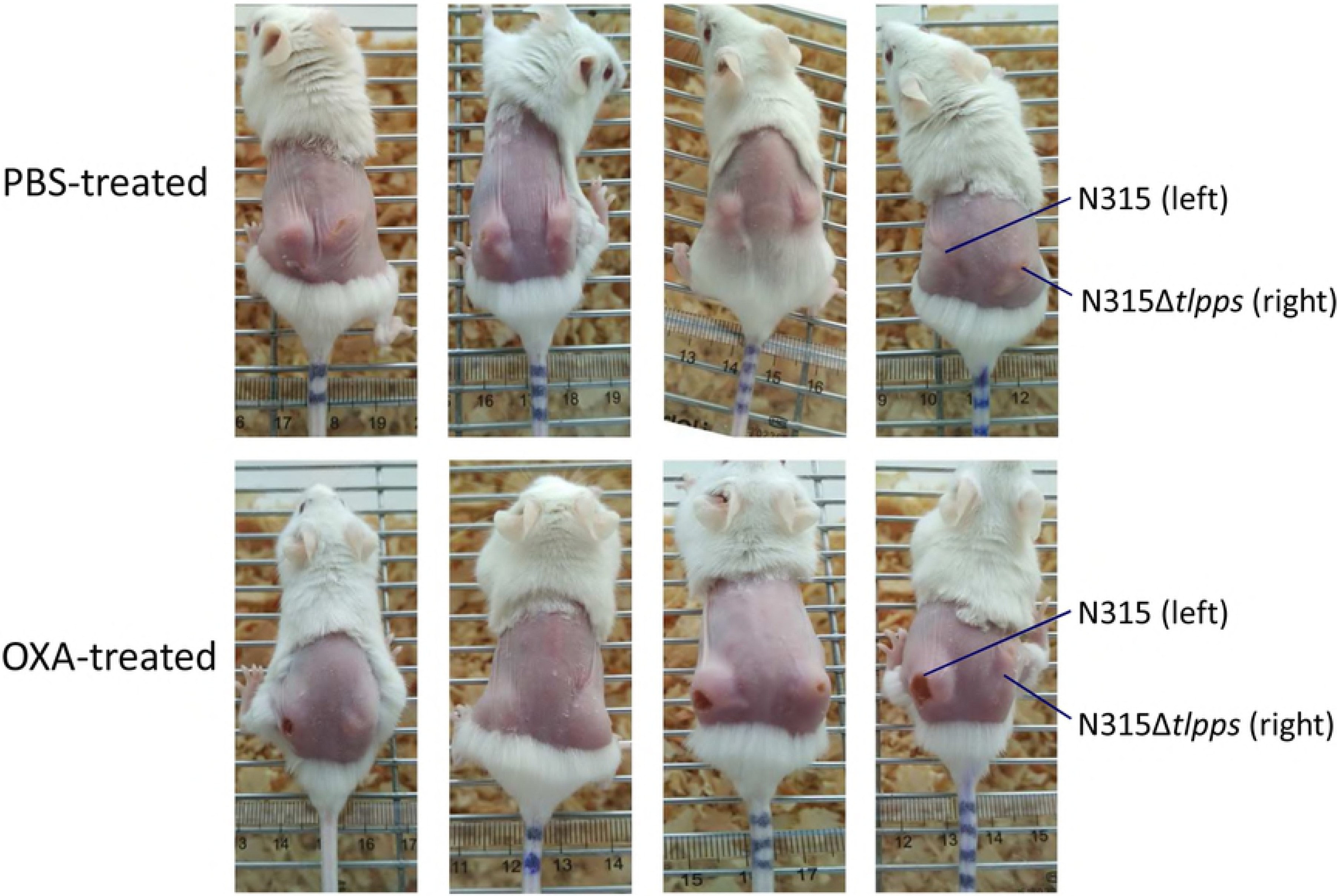
Abscesses in mice 7 days after infection. The mice were subcutaneously injected in both flanks with 5 × 10^7^ OXA-treated N315 and N315Δ*tlpps* cells and intraperitoneally injected with 1 μg of OXA per gram weight twice a day for 14 days. The skin lesions were recorded at the 7en day post-infection. The abscesses caused by the OXA-treated N315 were larger than those caused by the OXA-treated N315Δ*tlpps*, PBS-treated N315, and PBS-treated N315Δ*tlpps*.

**S6 Fig.**
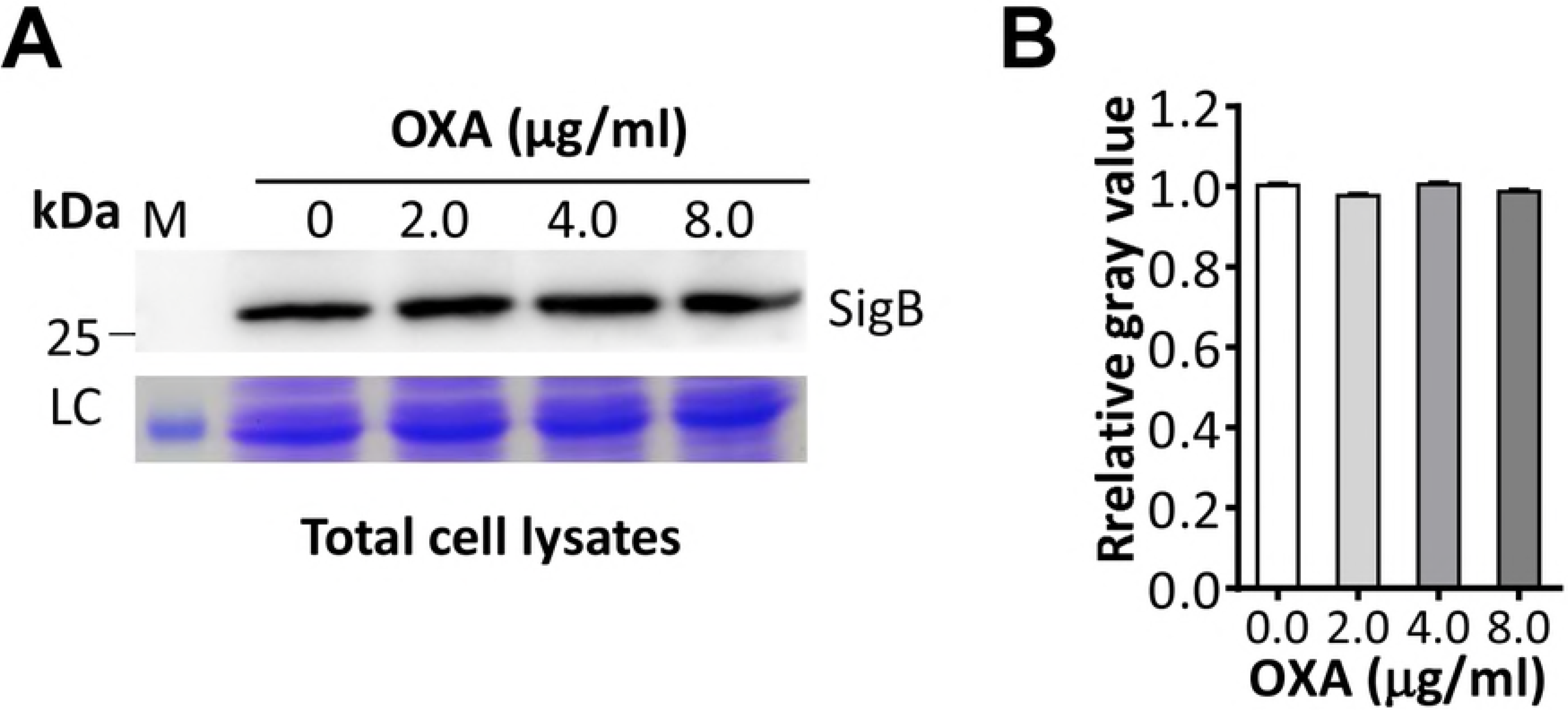
Western blot analysis of SigB in N315 with different concentrations of OXA treatment. (A) SigB was not changed in N315 with OXA treatment. The molecular weights of the protein marker (M) were indicated on the left. LC, loading control. (B) Evaluation of gray value of the N315 SigB in each lane using ImageJ software. The relative value of SigB/LC in the first lane (0 μg/ml OXA) was adjusted to 1.0, and the relative gray values in other lanes were calculated and indicated.

**S7 Fig.**
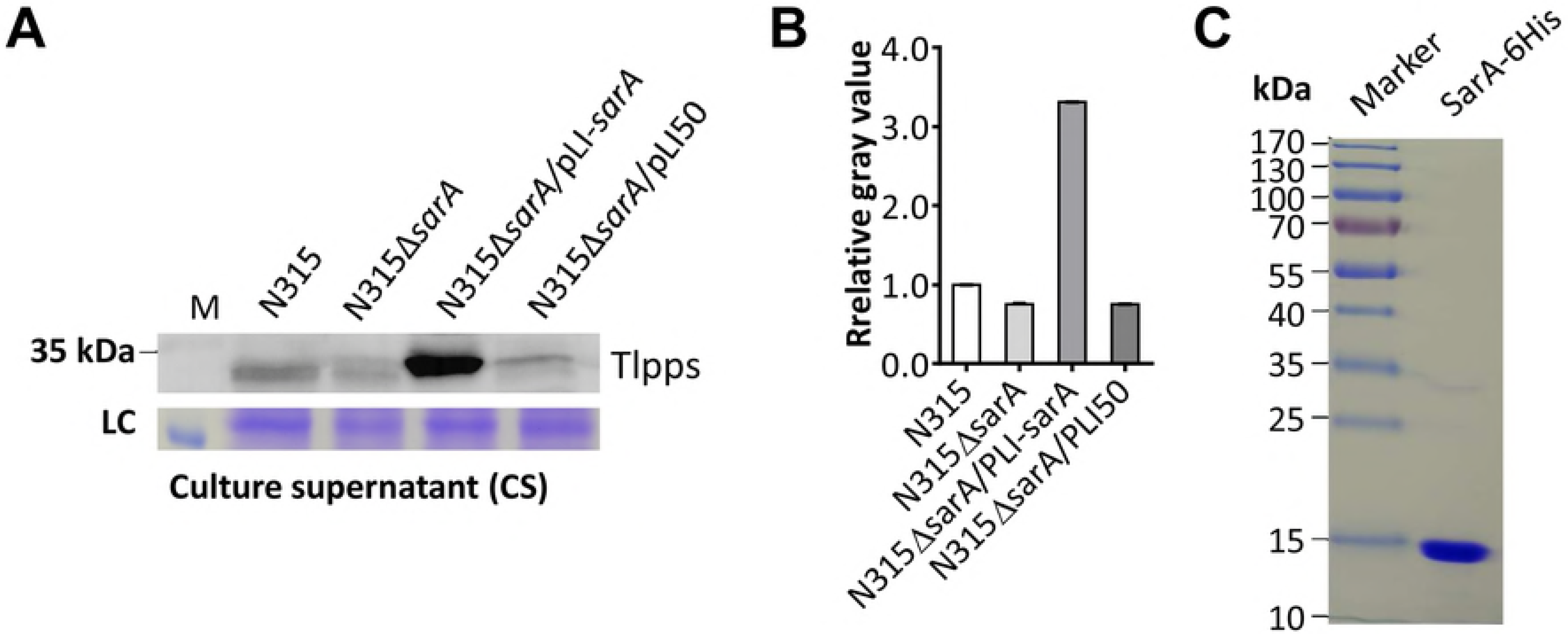
The deletion of *sarA* decreased the secretory Tlpp expression by N315. (A) Western blot analysis of the reduced expression of Tlpp proteins in the culture supernatant of N315Δ*tlpps*, which could be restored by the complementary *sarA* (N315Δ*sarA*/pLI-*sarA*) but not by the empty pLI50 plasmid-carrying strain (N315Δ*sarA*/pLI50). LC, loading control. (B) Evaluation of gray value of the Tlpp proteins in each lane using ImageJ software. The relative value of Tlpp/LC in the first lane (N315) was adjusted to 1.0, and the relative gray values in other lanes were calculated and indicated. (C) SDS-PAGE analysis of the recombinant SarA (His-tagged).

**S8 Fig.**
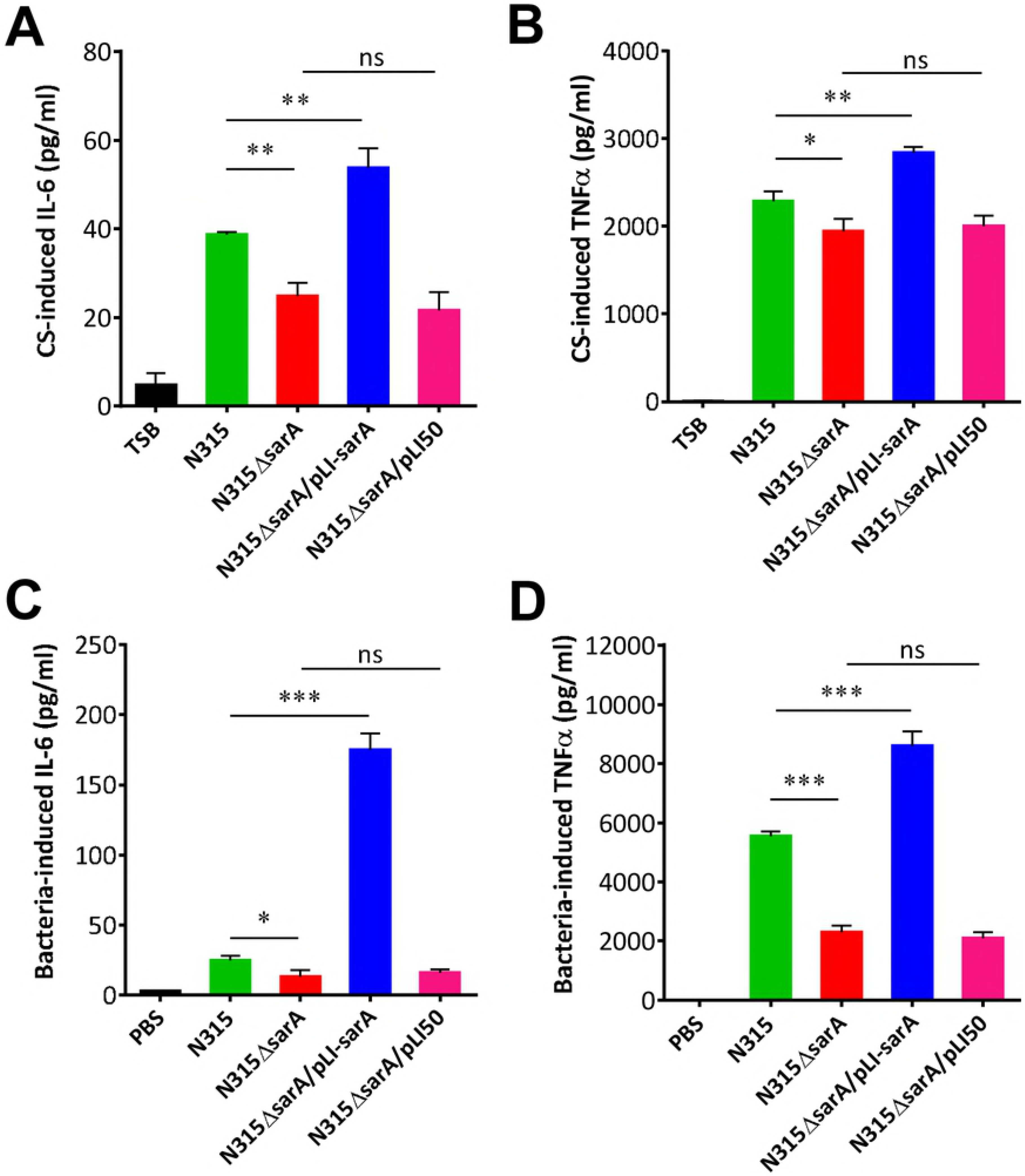
SarA mutant stimulated less IL-6 and TNFα production in macrophages. (A) IL-6 and (B) TNFα levels produced by macrophages post-treated with the culture supernatant of N315, N315Δ*sarA*, N315Δ*sarA*/pLI-*sarA*, and N315Δ*sarA*/pLI50, respectively. (C) IL-6 and (D) TNFα levels produced by macrophages infected with N315, N315Δ*sarA*, N315Δ*sarA*/pLI-*sarA*, and N315Δ*sarA*/pLI50 at an MOI of 30. The cytokine levels were determined by ELISA 6 h post-infection. The experiments in duplicate were conducted in thrice. ns represented no statistical significance, * *P* < 0.05, ** *P* < 0.01, *** *P* < 0.001.

**S9 Fig.**
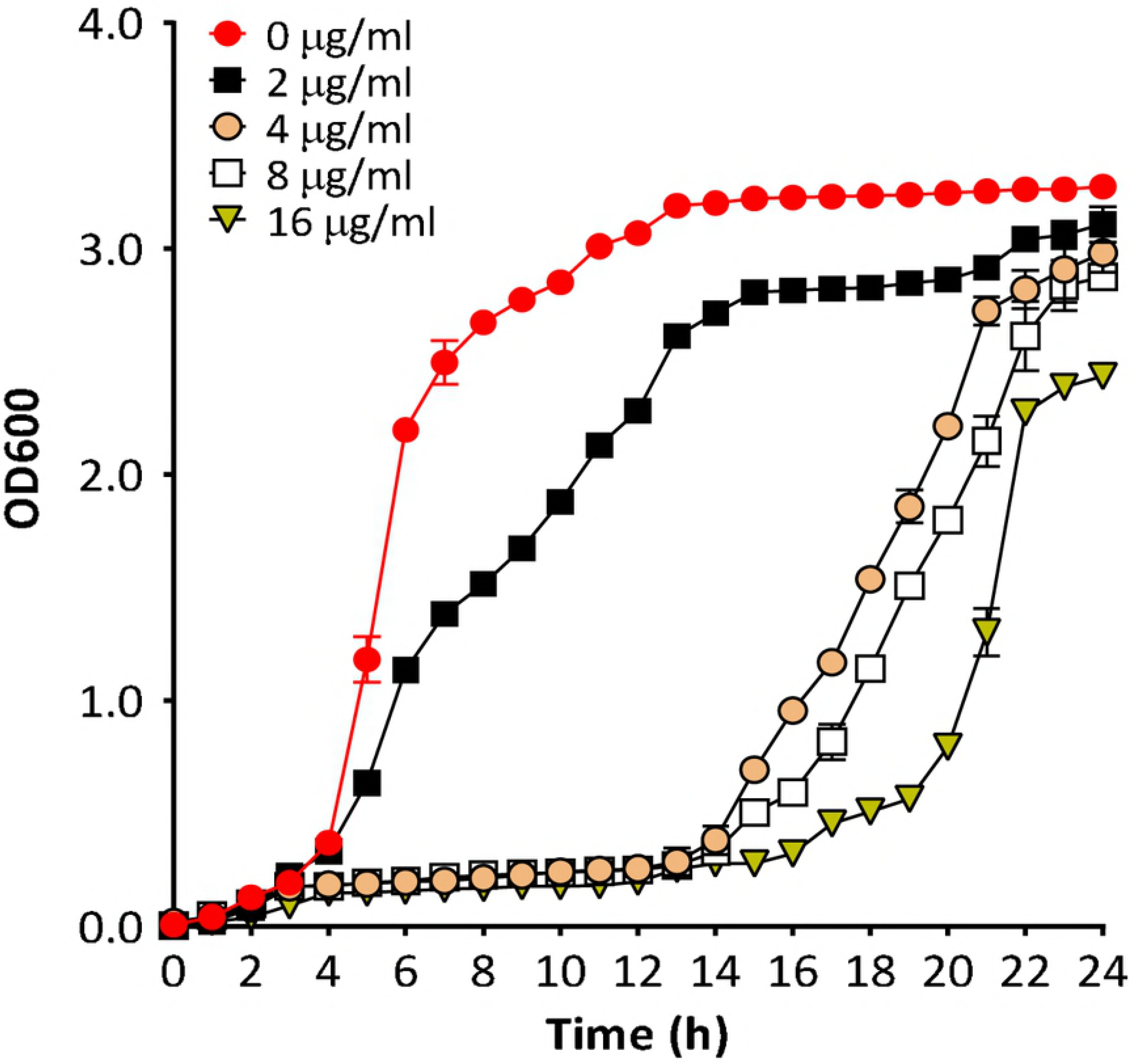
Growth curve of MRSA strain N315 in the culture media containing different concentrations of OXA as indicated. Independent growth rates of different concentrations of OXA treated N315 were determined. The experiment was conducted in thrice, and the data were shown as mean ± S.D. The growth of MRSA N315 was significantly inhibited by more than 4 μg/ml of OXA was administrated at the early stage, although the MIC value of N315 against OXA was higher as 512 μg/ml (S2 Table) because of the heterogeneous β-lactam-resistant phenotype of MRSA [46]. Therefore, the subinhibitory concentrations of OXA, 2 μg/ml, was used unless specifically stated in this study.

**S10 Fig.**
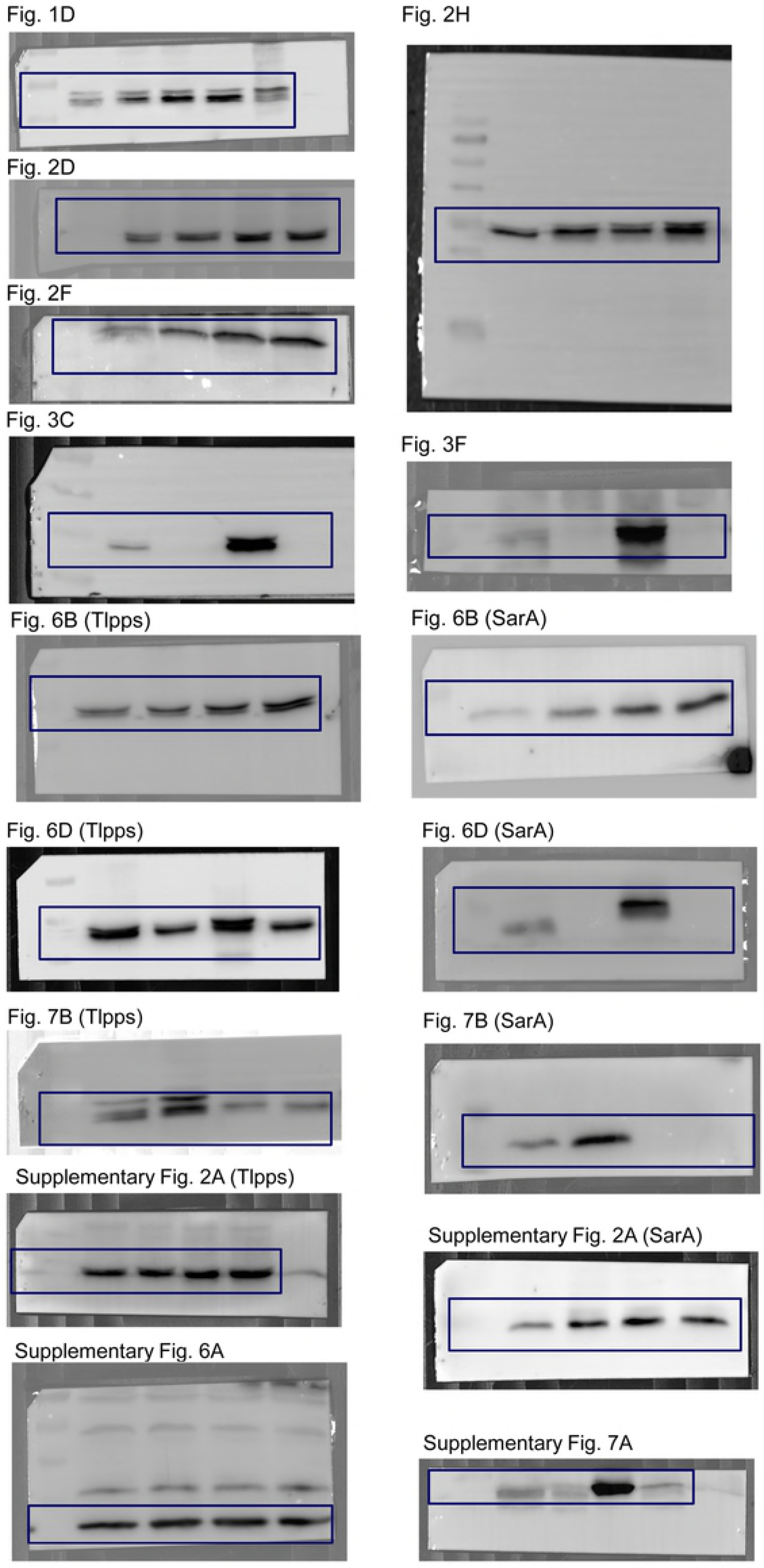
Full Western blot data. The full-length blots for all Western blot pictures. The black boxes represented the depicted parts of the blot.

**S1 Table. Predicted Lpps of *S. aureus* N315.**

**S2 Table. The MICs of MRSA strains.**

**S3 Table. Proteins identified in the protein band of β-lactams induced MRSA N315 by LC-MS/MS.**

**S4 Table. The distribution of** *tlpps* **cluster in major MRSA clones.**

**S5 Table. Primers used in this study.**

**S6 Table. Strains and plasmids used in this work.**

## Acknowledgments

We would like to thank the members of the Xiancai Rao and Xuhu Mao research groups for their critical reading of the manuscript. We thank Professor Baolin Sun (University of Science and Technology of China) providing plasmid pGFP and pLI50. We also thank Professor Lefu Lan (Shanghai Institute of Materia Medica, Chinese Academy of Sciences) for providing plasmid pOS1.

